# Precision phenotyping of a barley diversity set reveals distinct drought response strategies

**DOI:** 10.1101/2024.01.26.577377

**Authors:** Maitry Paul, Ahan Dalal, Marko Jääskeläinen, Menachem Moshelion, Alan H. Schulman

## Abstract

Plants exhibit a wide array of responses and adaptive mechanisms to drought. During drought, the trade-off between water loss and CO_2_ uptake for growth is mediated by the regulation of stomatal aperture in response to soil water content (SWC), among other factors. We earlier identified, in a few reference varieties of barley that differed by the SWC at which transpiration was curtailed, two divergent water use strategies: water-saving (“isohydric”) and water-spending (“anisohydric”). We proposed that an isohydric strategy may reduce risk from early droughts in climates where the probability of precipitation increases during the growing season, whereas an anisohydric strategy is consistent with environments having terminal droughts, or with those where dry periods are short and show little seasonal variation. Here, we have examined drought response in an 81-line barley diversity set that spans 20^th^ century European barley breeding and identified a several lines with a third, dynamic transpirational response to drought. We found a strong positive correlation between vigor and transpiration, the dynamic group being highest for both. However, these lines curtailed daily transpiration at a higher SWC than the isohydric group. While the dynamic lines, particularly cv Hydrogen and Baronesse, were not the most resilient in terms of restoring initial growth rates, their strong initial vigor and high return to initial transpiration rates meant that their growth nevertheless surpassed more resilient lines during recovery from drought. The results will be of use for defining barley physiological ideotypes suited to future climate scenarios.

## 1 Introduction

Drought is a ubiquitous abiotic stress that is increasing in frequency and severity as the amplitude of weather fluctuations grows due to climate change (Cohen et al., 2021, Seleiman et al., 2021, Yin et al., 2018). Plants respond to drought by reducing their evaporative water loss, concomitantly decreasing CO_2_ uptake for photosynthesis. The trade-off between water loss and CO_2_ uptake is mediated by the regulation of stomatal aperture in response to atmospheric CO_2_, ambient temperature, vapor pressure deficit, and light, as well as leaf hydration and soil water content (SWC) (Buckley, 2005, Kollist et al., 2014, Peters et al., 2018).

The susceptibility of an individual plant to drought stress and its ability to recover therefrom depends both on the length and intensity of the stress and on the adaptive capacity of the response mechanisms of the plant. The response mechanisms include signaling cascades that mediate stomatal closure (Merilo et al., 2015), as well as physiological and metabolic responses (Jogawat et al., 2021). Those include osmolyte accumulation for osmotic adjustment (Hildebrandt, 2018), enzymatic and non-enzymatic scavenging of excess reactive oxygen species (ROS) to mitigate dehydration and cellular damage (Das and Roychoudhury, 2014), and changes in the chloroplast proteome (Chen et al., 2021).

Under well-watered conditions, plants may differ in their water use efficiency (WUE), which is the carbon fixation or growth rate relative to the rate of transpirational water loss (Hatfield and Dold, 2019). Plants may be optimized for growth rate given non-limiting water, for water conservation, or for WUE. With the arrival of drought, two differing idealized strategies may be followed: isohydric or anisohydric (Moshelion et al., 2015, Sade et al., 2012). Isohydricity implies closing of stomata at a relatively high SWC to maintain a relatively constant water potential, thereby sacrificing carbon fixation but delaying plant dehydration, and comprises many physiological parameters (Scharwies and Dinneny, 2019). In contrast, plants with anisohydric behavior keep their stomata open to a relatively low SWC, allowing the leaf water potential to decline (Hoshika et al., 2020, Negin and Moshelion, 2016). The terms are often used, as here, loosely to refer to water use strategy, respectively water-conserving (isohydric) and non-conserving (anisohydric), rather than referring to the actual hydric status of the leaf, due to the practical difficulty in measuring leaf water potential non-destructively during the course of a drought experiment (De Swaef et al., 2022, Sade et al., 2012).

Barley is the world’s fourth most widely cultivated cereal and is grown on every continent (FAOSTAT, 2023, Lister et al., 2018). It is either cultivated as a spring crop, which is sown in the spring and harvested in the late summer autumn or as a winter crop, which is sown in the autumn or early winter and harvested in the spring (Lister et al., 2018). Normally, spring-sown barley will experience periodic droughts during the growing season, whereas winter-grown barley will encounter terminal drought during grain filling and maturation (Hakala et al., 2012). In an earlier study of several reference varieties of barley that differed by the SWC at which transpiration declined (Paul et al., 2023), we proposed that an isohydric strategy may reduce risk from early droughts in climates where the probability of precipitation increases during the growing season (Paul et al., 2023), whereas an anisohydric strategy is consistent with environments having terminal droughts, or with those where dry periods are short and show little seasonal variation. In recent analyses of four high-yielding European spring barley cultivars subjected to a standardized drought treatment imposed around flowering time we found, moreover, that one variety (RGT Planet) displayed a dynamic drought response (Appiah et al., 2023). This variety displayed high transpiration under ample water supply, but switched to a water-conserving phenotype upon drought.

Here, we have examined drought response in an 81-line barley diversity set, which spans (Fig. 1B, Table S1) 20^th^ century European barley breeding. A high-precision lysimeter platform (Dalal et al., 2020) permitted highly regulated irrigation and continuous monitoring of soil and plant water relations, plant stomatal response, and biomass increase. We looked at differences in rates of transpiration and growth under well-watered conditions, transpirational responses to SWC during drought, and the degree of recovery following rewatering. We found that a dynamic transpirational response to drought is not unique to RGT Planet but represents a third strategy among barley cultivars. The results will be of use for defining barley physiological ideotypes suited to future climate scenarios.

**Figure 1.**
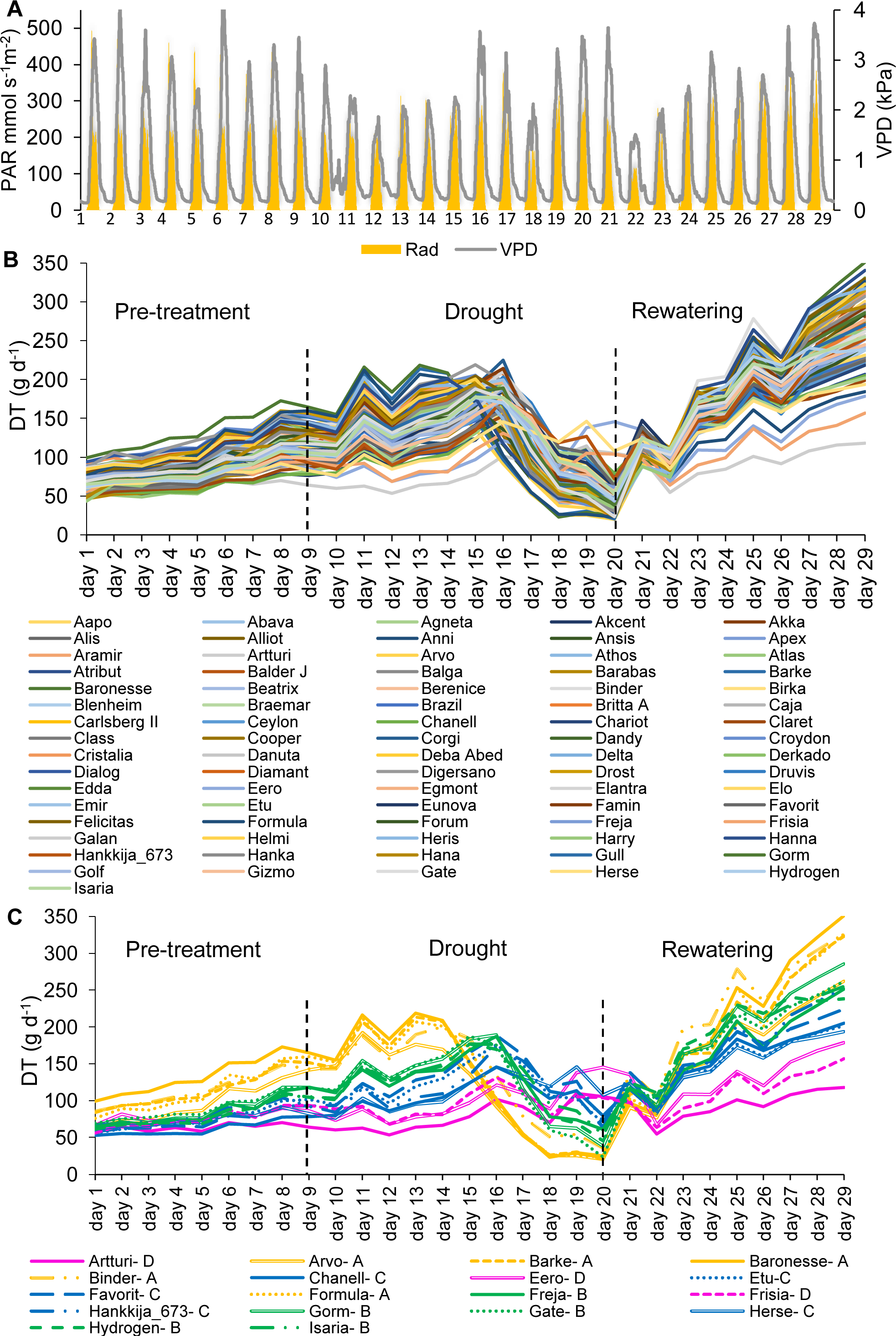
Daily Transpiration (DT) of the 81-line barley population grown on the lysimeter system for first screening. (A) Daily vapor pressure deficit (VPD) and Photosynthetic Active Radiation (PAR) during 29 consecutive days of the experiment. (B) DT in response to the soil-atmosphere water gradient. Each line represents a single plant for each accession. (C) DT of 18 selected lines within the 81-accession experiment during the pre-treatment, drought and rewatering shown in (B). Four groups of lines were identified in the 81-line experiment, based on their DT and stomatal closure. These are labeled as A (yellow), B (green), C (blue), D (magenta). Dashed black lines at Day 9 and Day 20 show the beginning and end of drought treatment.

## 2 Materials and methods

### 2.1 Plant material

The barley lines used in this screening experiment were from a diversity set assembled to represent the breadth of European barley breeding during the 20^th^ century (Xu et al., 2018). The set studied here comprised 81 spring barley lines, 72 two-row and 9 six-row, from 14 countries. Information on the characteristics of 18 barley lines, including their release year, country of origin, pedigree and breeders are detailed in Tables S1 and S2. A subset of 18 varieties were selected from the 81 as exemplars of their water use strategy, as described below. The subset comprised 12 two-row and 6 six-row lines from 7 countries. The 18 chosen lines are distributed across the diversity space (Figure S1), as revealed by 864 gene-based single nucleotide polymorphism (SNP) markers (Tondelli et al., 2013).

### 2.2. Growth conditions

Seeds were sown into 50 ml cones filed with peat soil, on trays, one seed per cone, and the trays covered with plastic wrap and aluminum foil for two weeks at 4°C as a means to break dormancy and to enhance germination (Galkin et al., 2018). The trays were then placed in a controlled glasshouse under short-day conditions (8/16 hr light/dark, 16/10°C day/night). Following emergence, the seedlings were grown for either 8 weeks (81-line experiment) or 12 weeks (18-line experiment) in total. Seedling roots were washed and the seedlings transplanted into potting soil (Dalal et al., 2020, Paul et al., 2023) in 4L pots and placed into a semi-controlled greenhouse. After two weeks of acclimatization, the pots were mounted on the lysimeter system (Figure S2). To maximize homogeneous exposure to the ambient conditions on the lysimeter platform, all the pots were placed in random order. Temperature and relative humidity (RH) were respectively around 24-33°C and 30-60% for the 81-line experiment, and 20-35°C and 20-80% for the 18-line experiment.

The PlantArray lysimeter platform (Plant-Ditech Ltd., Israel) (Dalal et al., 2019) includes a meteorological station and continuously records the physiological conditions of the experiment and is equipped with an automated irrigation system. During the 81-line and 18-line experiments, the system recorded minimum and maximum mid-day daily light intensities ranging from 116 to 493 mmol s^-1^ m^-2^ and 74 to 514 mmol s^-1^ m^-2^ respectively (Figures 1A, S3A), and vapor pressure deficits (VPD) of 1.5 to 4.3 kPa (average 2.6 kPa) and 0.6 to 4.8 kPa (average 2.2 kPa) respectively.

### 2.3 Experimental setup

Drought experiments were carried out as previously on the PlantArray platform, which is a high-throughput physiological diagnostic system consisting of a highly sensitive, temperature compensated multi-lysimeter array (Appiah et al., 2023, Dalal et al., 2019, Halperin et al., 2017, Paul et al., 2023) and comprised three phases: pre-treatment, drought, and rewatering. In pre-treatment, the plants were maintained on the lysimeter platform and were well watered. Under drought, the plants were exposed to dehydration by limiting the water content in each pot until they reached a specific SWC. During rewatering, plants were irrigated as in the pre-treatment phase. The 81-line experiment lasted a total of 29 days: 9 days of pre-treatment, 11 days of drought, and 9 days of rewatering. For the subsequent 18-line in-depth evaluation, the set was given 11 days of pre-treatment (Figure S2B), followed by 15 days of drought (Figure S2C) to attain a similar SWC as in the first set. The drought period was terminated for each line individually as soon as it fell below 15% SWC. All lines were then given 29 days of rewatering for recovery (Figure S2D). Due to the second experiment’s average atmospheric VPD being lower than that of the first, the drought treatment was longer. The total elapsed time of the second experiment was 55 days.

### 2.4 Measurements of physiological traits

The PlantArray platform continuously collects data and sends it to a central computer for additional analysis by SPAC (soil–plant–atmosphere continuum) analytical software of soil and atmosphere data alongside plant traits (Dalal et al., 2020). The processed data allowed us to determine the physiological parameters of single whole plants in individual pots simultaneously for the entire experimental time, including growth rate, transpiration, stomatal conductance, and WUE (Dalal et al., 2019, Halperin et al., 2017). The initial 81-line screening was carried out with only one biological replicate. The subsequent 18-line experiment was made in five to eight replicates. We chose five consecutive days during each of the pre-treatment, drought, and rewatering stages, where the VPD and PAR were most stable, to analyze the quantitative physiological traits (QPTs) of the lines in detail. Over the course of the 18-line experiment, these periods comprised days 7 to 11 during pre-treatment, 22 to 26 during drought, and days 51 to 55 during recovery. Using SPSS software, means and standard errors were determined and the one-way ANOVA Tukey Post-Hoc test at a significance level of *p*=0.05 was carried out for multiple comparisons of the lines. In the reported analyses, lines without significant differences share superscript letters.

## 3 Results

Altogether 81 lines were screened without replication for drought stress response and yield-related QPTs. Of this set, 18 were then chosen for more intensive investigation. For the 18 lines, 5 to 8 biological replications were made, creating a total of 139 plants. For both sets of experiments, a well-watered period, with irrigation at night until each pot reached its full capacity, was followed by a drought treatment in which irrigation was minimized. The QPTs were monitored in real time on a multi-lysimeter platform, with data collected every few minutes by SPAC analytical software throughout the experimental period.

### 3.1 Water use strategies during drought and recovery

*Daily transpiration*: The VPD and PAR during the whole experimental period is shown in Figure 1A. During the well-watered pre-treatment, daily transpiration of all 81 barley lines increased in parallel but diverged as the plants grew at different rates (Figure 1B). During the drought, the lines responded differentially regarding the day and degree to which their daily transpiration (DT) dropped. Similarly, following rewatering on day 21, though all the plants’ DT grew substantially throughout the rewatering phase, the lines responded disparately to water availability and showed varying degrees of recovery rate - resilience. The 81 barley lines could be divided into four groups by their transpirational behavior, from which we extracted 18 lines (Groups A, five lines; B, five lines; C, five lines; D,3 lines) as representative of the range (Figure 1C) for the second set of experiments.

*Physiological drought point* (θ_c_): Theta-crit (θ_c_) is the critical SWC (The point at which the soil water content becomes a limiting threshold for supporting maximal transpiration values) at which the plant responds by reducing its transpiration rate (Dalal et al., 2019, Halperin et al., 2017), shown in Figure 2A for the 18 chosen barley lines during the 81-line experiment. The four groups (Figure 1C) define four distinct patterns for combinations of θ_c_ and the maximum transpiration rate (TR_max_) under well-watered conditions. Group A and B had similar θ_c_, at about ∼30% SWC, though at different DT; Group C had a still lower DT and a very low θ_c_, at ∼20% SWC; Group D had the highest θ_c_, but the lowest DT. Comparison between the TR_max_ and θ_c_ under drought (Figure 2B) shows a significant difference between each group and a correlation between the two measures, TR_max_ declining from Group A successively to B, C, and D under well-watered conditions. The groups showed distinct rates of decline in transpiration rate vs SWC, with Groups A and B declining slowly and Groups C and D precipitously (Figure 2A). Table S3 contains TR_max_ and θ_c_ for each of the 18 line subset.

**Figure 2.**
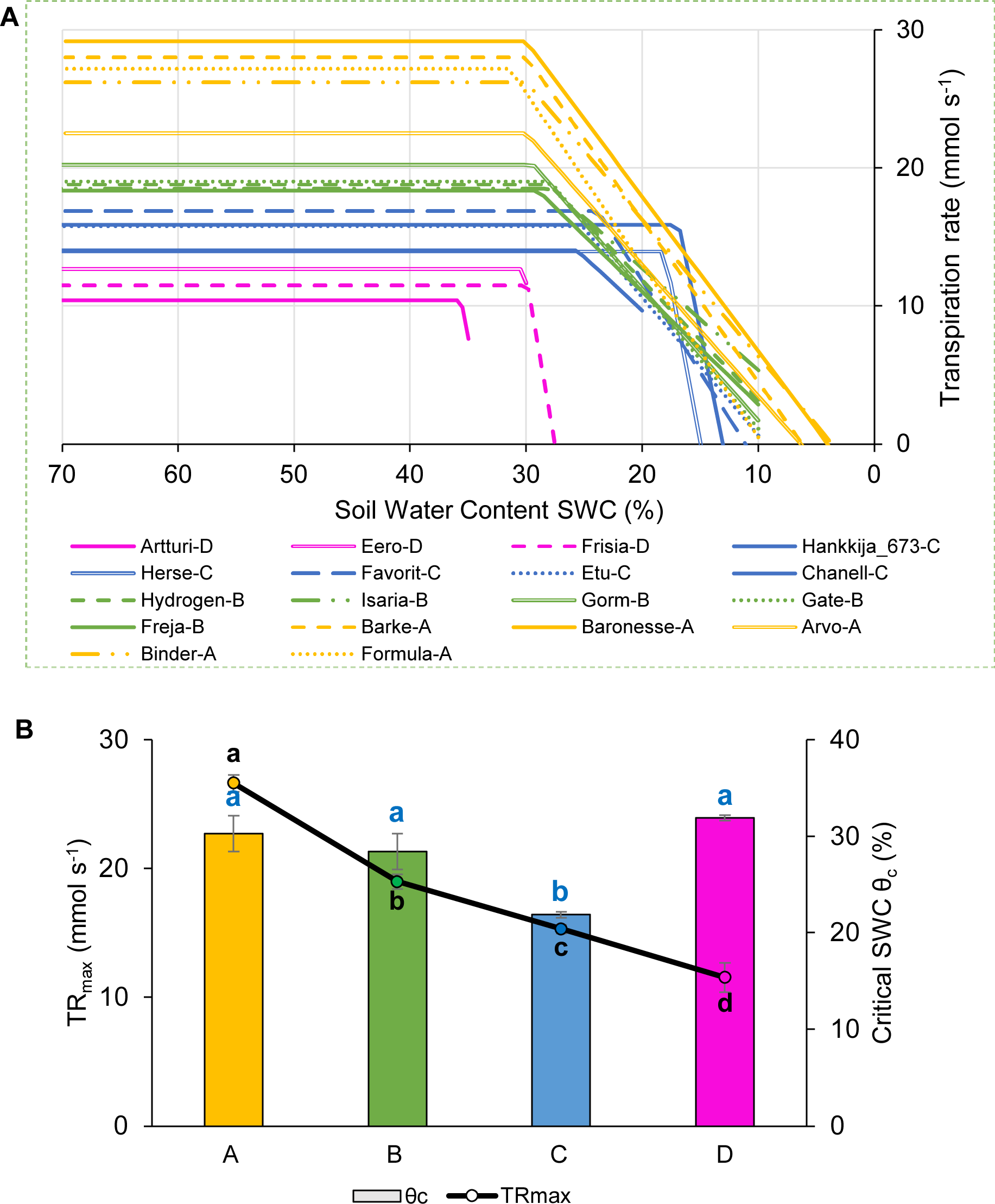
Transpiration rate response to drought during 81-line screening. Behavior of 18-line subset shown. (A) Piece-wise fitted line between midday whole plant transpiration rate (mmol s^-1^) and soil water content (SWC). The four physiological groups are as in Figure 1: A (yellow), B (green), C (blue), D (magenta). (B) Maximum transpiration rate (TR_max_) and θ_c_ averaged for the four groups. Lowercase letters, significance levels for TR_max_ (black) and θ_c_ (blue).

### 3.2 Whole-plant water relations during drought and recovery

In the replicated and more extensively analyzed experiments with the 18-line subset, we regrouped the lines to take into account their behavior through drought and recovery. Group 1, with highest transpiration, had four lines (Hydrogen, Hankkija 673, Baronesse, Isaria), followed by Group 2 (Etu, Gorm, Gate, Frisia, Barke, Chanell, Favorit) and Group 3 (Arvo, Herse, Formula, Artturi, Eero, Binder, Freja) with seven lines each (Figure 3). During pre-treatment, the overall transpiration of all barley lines ranged from 73 to 207 g day^-1^ (Figure 3A), which dropped to 36 to 131 g day^-1^ (Figure 3B) during drought stress and recovered to 174 to 380 g day^-1^ (Figure 3C). The maximum DT of 207 g day^-1^ during a pre-treatment was observed in Hydrogen, while the lowest DT was observed in Freja at 73 g day^-1^.

**Figure 3:**
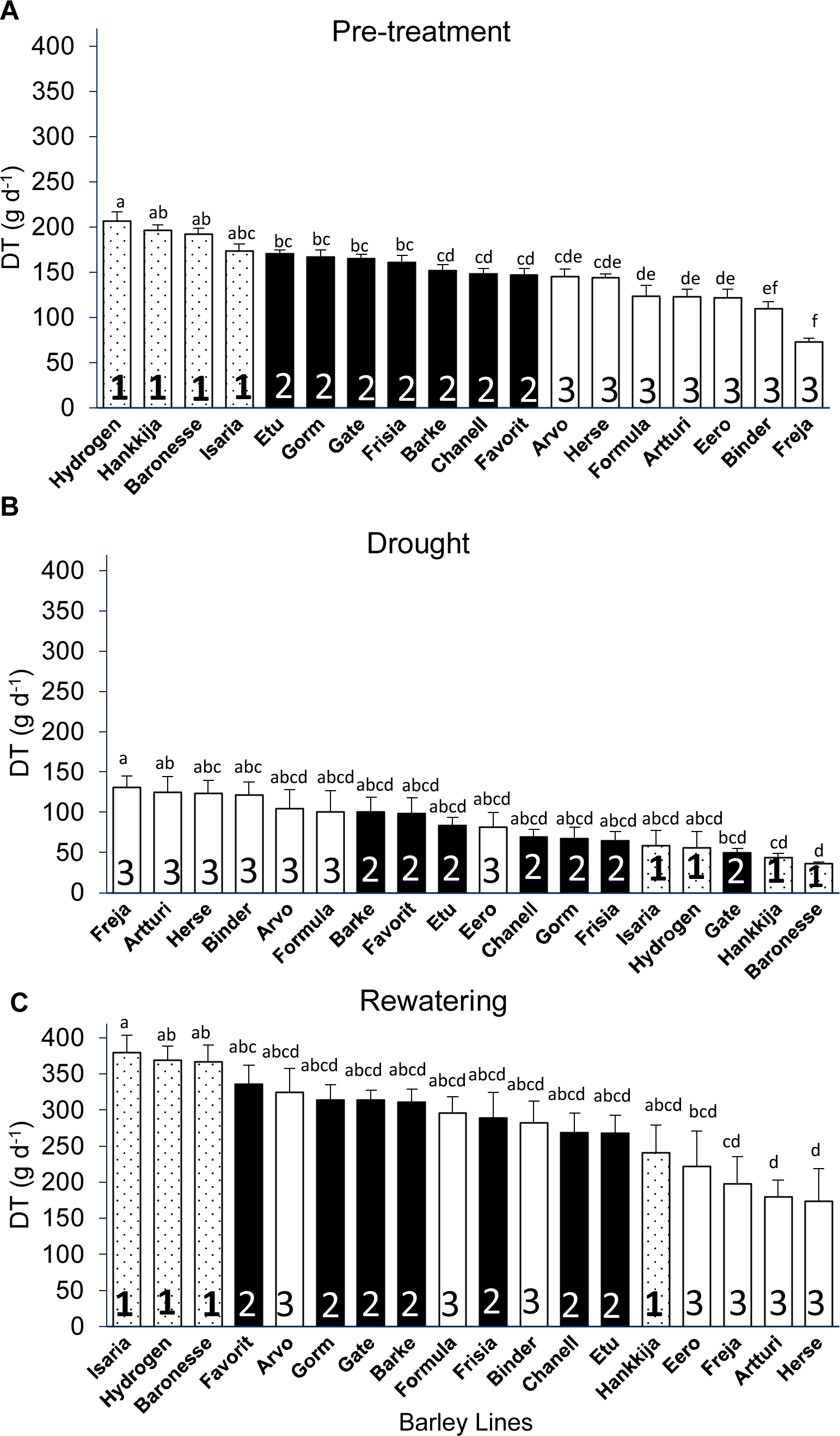
Daily Transpiration (DT) of 18 barley lines grouped by significance levels. Each treatment is the average of DT from 5 consecutive days when the PAR and VPD were the most identical, and for each line 5 to 8 biological replicates were measured. (A) Pre-treatment (well-watered); grouping from highest to lowest DT (average of day 7-11). (B) Drought DT, lines arranged in descending order (average of day 22-26). (C) Rewatering DT, lines arranged in descending order (average of day 51-55). Lowercase letters above the bars show significant difference between lines, those with no significant difference receiving the same letter. Group 1 has the highest DT, followed by Group 2 and Group 3. Tukey’s Post Hoc multiple comparisons test done in SPSS, with bars representing mean ± SE.

During drought, we observed Group 1 transition from the highest to one of the lowest DT and Group 3 shift from lowest to highest, with the exception of Eero, which remained in the center of the range. The majority of the lines in all three groups reverted to their pre-treatment positions throughout the rewatering phase. Hankkija 673, Arvo, and Formula, on the other hand, responded differently during rewatering, remaining at a transpiration level similar to that during drought, not returning to the pre-drought level.

Except for a few outliers, most of the lines with intermediate pre-treatment DT (Group 2) remained in the center of the distribution during both the drought and the recovery. When the lines were compared between the treatments, we found a strong negative correlation between pre-treatment and drought *r* = -0.83 (Figure S4A), which is driven by the opposing behaviors of Groups 1 and 3.

There was a weak positive correlation, *r* = 0.58 between pre-treatment and rewatering (Figure S4B), and a weak negative correlation for drought versus rewatering *r* = -0.59 (Figure S4C), likewise driven by differential behaviors by Groups 1 and 3. The daily transpiration of the 18 barley lines with all biological replicates (139 plants) for 55 consecutive days during pre-treatment, drought, and rewatering (Figure S3B), with daily fluctuation in VPD and PAR (Figure S3A), shows similar variations as found in the first, 81-line screening (Figure 1B, 1C).

Transition to drought response at θ_c_ was highly correlated (*r*=0.89) with whole plant transpiration levels (E_max_) (Figure S5B); three lines from Group 1 (Hydrogen, Baronesse, Hankkija 673) had both the highest SWC at θ_c_ and the highest E_max_ (Figure 4). Freja had both the lowest transpiration (E_max_ =8.8 ± 0.8 mmol s^-1^g^-1^) and θ_c_ (26.7 ± 1.2 %). All the lines fell within a transpiration range of 8 to 15 mmol s^-1^g^-1^, with all θ_c_ values between 26 to 42 % SWC (Figure 4, Table S4). Hydrogen had not only the highest transpiration (E_max_ = 15.1 ± 1.3 mmol s^-1^g^-1^) but also the highest θ_c_ (42.6 ± 2.9 %).

**Figure 4.**
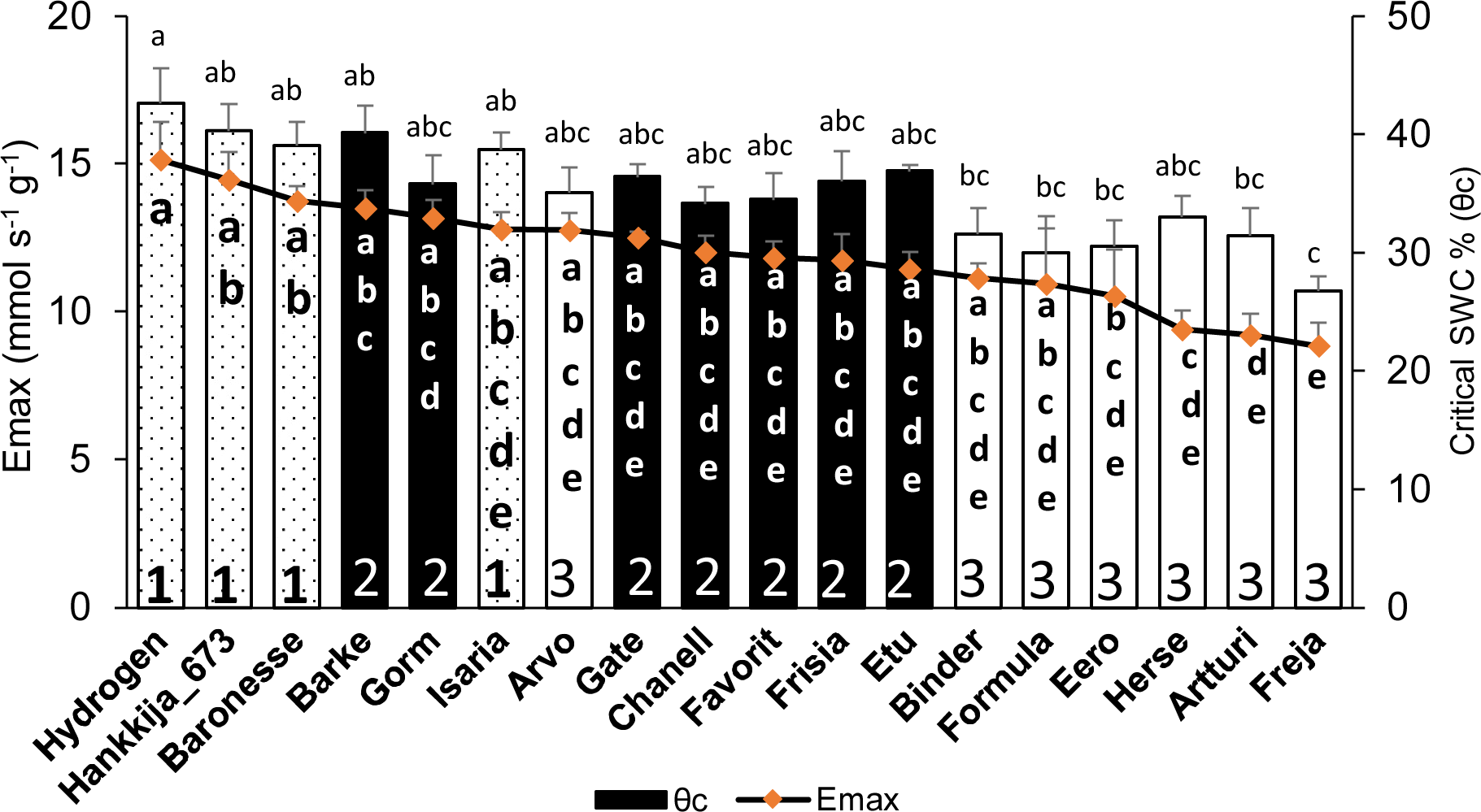
Comparison between midday whole plant transpiration (Emax) during pre-treatment and soil water content (SWC) at θ_c_ in 18 barley lines. Lowercase letters inside the bars are significance groups for Emax; those above the bars are for θ_c_.

*Canopy Stomatal conductance (GSc):* During pre-treatment, Hydrogen (816 mmol s^-1^g^-1^) had the highest GSc, whereas Freja (428 mmol s^-1^g^-1^) had the lowest (Figure 5A). The DT and GSc are highly (*r=*0.94) correlated, with Groups 1,2, and 3 showing differential response (Figure S5A). When compared by DT group during pre-treatment (Figure 3A), the three lines from Group 1 with the greatest GSc (Hydrogen, Hankkija 673, Baronesse) also had the highest DT, except Isaria, which was in the middle. Similarly, the six lines from Group 3 (Freja, Artturi, Binder, Herse, Eero, and Formula) had the lowest GSc (Figure 5A) and DT (Figure 3A) values, apart from Arvo in the middle. Nevertheless, the majority of the lines in GSc pre-treatment Group 2 (filled bars) are in the same location as the DT pre-treatment.

**Figure 5:**
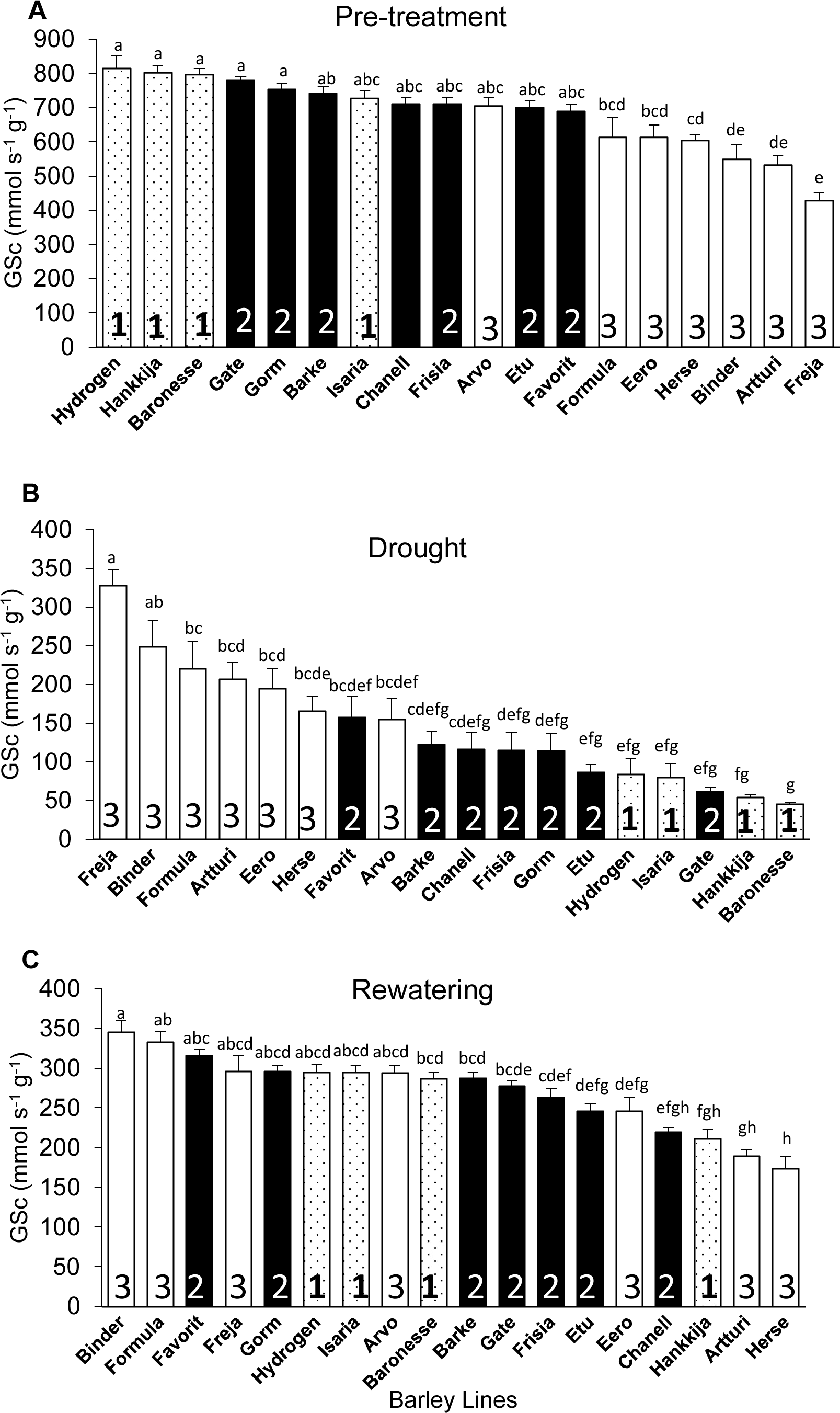
Canopy Stomatal Conductance (GSc) of 18 barley lines grouped by DT during pre-treatment. Each treatment is the average of GSc from 5 consecutive days when the PAR and VPD were most uniform; each line comprises 5 to 8 biological replicates. (A) Pre-treatment; lines arranged by descending GSc (average for days 7-11). (B) Drought; lines arranged by descending GSc (average for days 22-26). (C) Rewatering; lines arranged by descending GSc (average for days 51-55). Lowercase letters indicate significance groups. Bars represent mean ± SE.

During drought, however, Freja (highest) had 7.2X greater GSc than Baronesse (lowest) (Figure 5B), although the difference between the highest and lowest GSc was only 2X during pre-treatment and rewatering (Figure 5A, C). Like for DT, we observed a shift in Group 3 from lowest to highest GSc under drought, and a flip from highest to lowest GSc in Group 1 (Figure 5B). This was also evident when GSc during pre -treatment was compared to GSc during drought using Pearson correlation (*r*), which revealed a negative correlation of 0.95 (Figure S6A).

During rewatering (Figure 5C), we saw a distinctive GSc pattern in which only three lines (Herse, Artturi, and Eero) out of seven in Group 3 returned to their pre-treatment behavior, while the other three (Binder, Formula, and Freja) did not shift, showing no resilient behavior. Interestingly, Arvo remained in the middle group through all three experimental phases. From Group 1, three lines (Hydrogen, Isaria, and Baronesse) moderately recovered GSc, but not Hankkija 673, which recovered poorly, as it did for DT. Correlation analysis also supports the GSc pattern, where only 2% (*r* = 0.02) of the lines recovered from stress after 29 days of rewatering (Figure S6B), whereas 58% of the lines showed recovery in DT (Figure S4B).

*Calculated Plant Weight Gain (CPW):* Once rewatering relieves drought-induced desiccation, calculated plant weight (CPW) gain per day indicates resumption of growth and recovery. Like for DT and GSc, CPW for Hydrogen (22 g day-1) is significantly higher than for Freja (6 g day-1; Figure 6A). Group shifts between experimental phases (Figure 6A, B, C) are similar to those for DT (Figure 3A, B, C). Pre-treatment and drought CPWs are negatively correlated (*r* =-0.68; Figure S7A), whereas pre-treatment and recovery show a positive correlation (*r* = 0.68; Figure S7B). Altogether 68 % of the 18 lines recovered from stress. Comparing drought and recovery, we found that line ranking shifted in a similar way for DT and CPW with respect to GSc.

**Figure 6:**
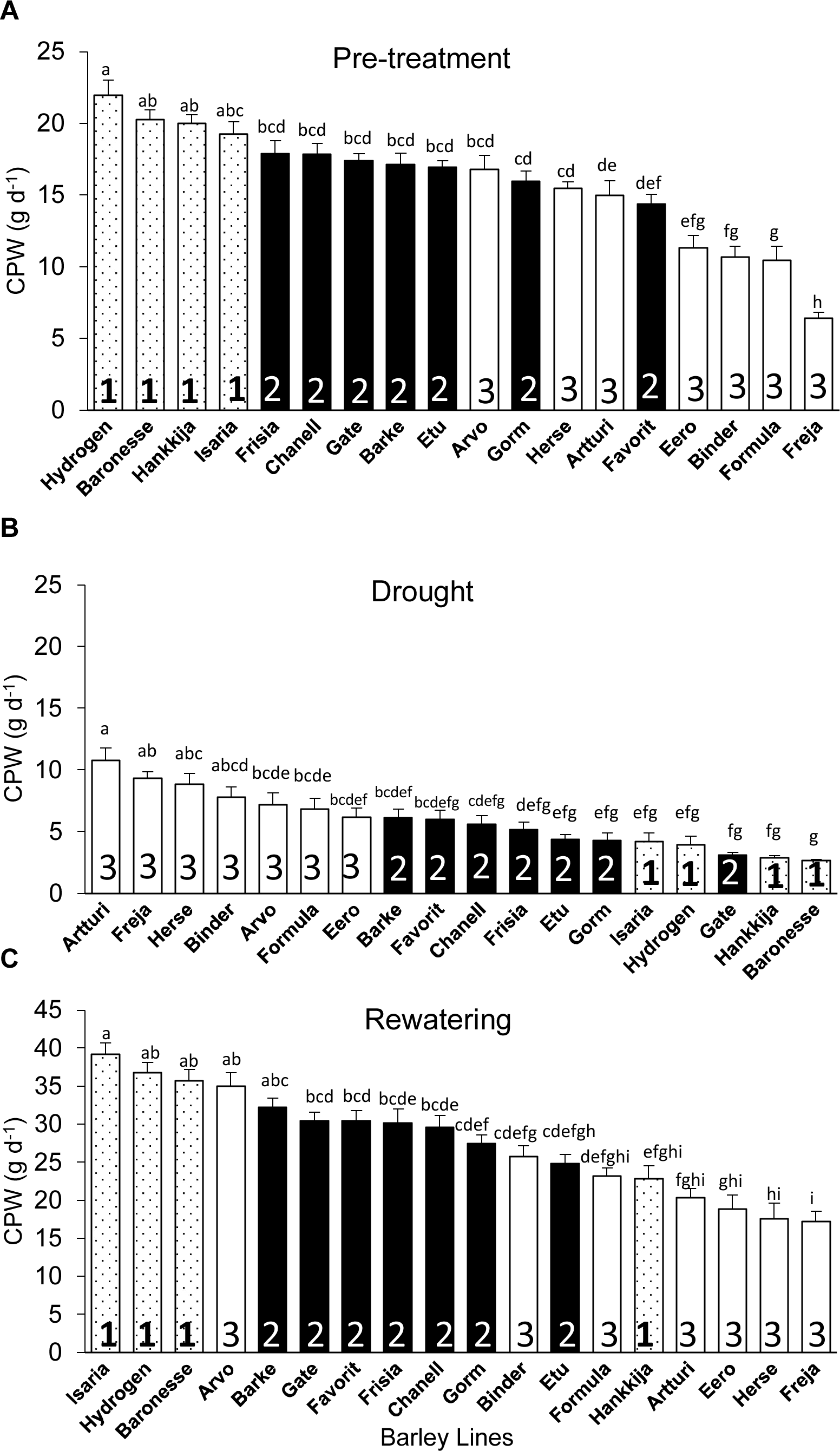
Calculated Plant Weight (CPW) gain per day for the 18 barley lines grouped based on significant levels during DT-pre-treatment. Each treatment is the average of CPW from 5 consecutive days when the PAR and VPD are the most identical, and each line is from of 5 to 8 biological replicates. (A) Pre-treatment (average of day 7-11). (B) Drought (average of day 22-26). (C) Recovery (average of day 51-55). For each phase, lines are arranged in descending CPW order. Lowercase letters indicate significance groups. Bars represent mean ± SE.

### 3.3 Plant vigor and the impact of stress

The line groupings (1, 2, 3) based on DT proved meaningful also for vigor, defined as average gain in CPW per day during pre-treatment, with the Group 1 lines showed the highest vigor (Figure 7A). Most of the lines in Group 3 had the lowest vigor, Freja being the poorest. However, during drought, Group 3 had the highest daily CPW (Figure 6B). If resilience is considered a return to a pre-drought CPW rate, then three Group 3 lines showing the highest resilience, but three within Group 3 the lowest (Figure 7B). However, Group 3 lines, which have a high (or non-responsive) CPW during drought (Figure 6B), returns to a low daily CPW in absolute terms (Figure 6C). The lines displayed as well differential recovery of DT during rewatering compared with drought conditions (Figure 7C). The high recovery set included: Hydrogen and Baronesse, comprising half of Group 1; Gorm, the only line from Group 2; Freja, Formula, and Binder of the seven lines in Group 3.

**Figure 7.**
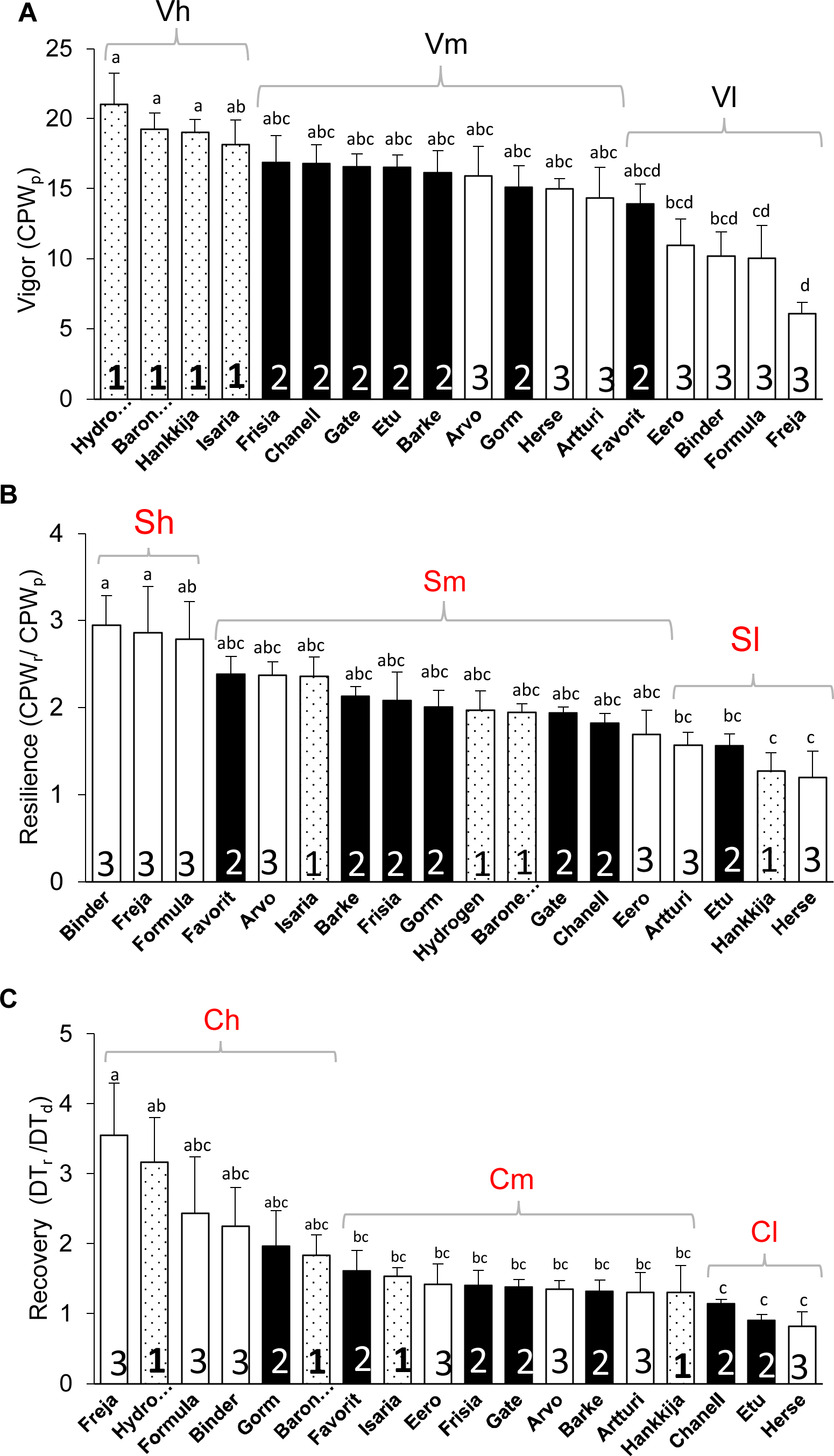
Vigor, Resilience, and Recovery of 18 barley lines. (A) Vigor, the average CPW from day 7 to 11 during pre-treatment (CPW_p_). (B) Resilience, the ratio between average CPW from day 51 to 55 during rewatering (CPW_r_) and pre-treatment CPW (CPW_p_), day 7 to 11. (C) Recovery, the ratio between average DT during rewatering, day 51 to 54 (DT_r_) and drought, day 18 to 26 (DT_d_). Means ± SE are displayed; lowercase letters indicate significance groups. Sets in brackets based on significance (lowercase letters) are high (h) medium (m), and low (l) Vigor (V), Resilience (S), and Recovery (C). The lines under high, medium and low vigor, resilience and recovery are presented in Table S5.

### 3.4 Water use efficiency and yield

The quantity of biomass or grain produced per unit of water transpired gives an overall idea of the tradeoffs between carbon fixation and water transpiration made during the life cycle of the plant. First, we calculated the mean total biomass (combined dry weight of husk, seed, and shoot; Figure S8A), yield (Figure S8B), and seed number (Figure S8C) from each biological replicate for all barley lines. Biomass, yield, and seed number varied respectively from 31.7 to 78.3 g, 0.4 to 14.3 g, and 9.6 to 380.4 per replicate. Isaria (78.3 ± 3.8 g) produced the highest biomass but one of the lowest yields (0.61 ± 0.06 g) and seed numbers (39 ± 5.9), hence, lowest harvest index (0.69 ± 0.08 %). Eero had the lowest biomass (31.7 ± 5.1 g), a low yield (1.78 ± 0.5 g) and moderate seed number (54.8 ± 20.2). Hydrogen had high total biomass (72.2 ± 3.5 g) but poor yield (1.3 ± 0.2 g) following drought; Freja produced low biomass (41.3 ± 5.6 g) and yield (0.4 ± 0.1 g) as well as the lowest seed number (9.6 ± 2.7). Notably, Hankkija 673, which is a six-row spring barley from Finland released in 1973, had a modest biomass output (60.3 ± 3.9 g) but one of the highest yields (10.2 ± 1.4 g) and seed numbers (380.4 ± 37.1). As a six-row variety, Hankkija had only a middling seed weight (TGW; Figure S9). The high yield of Herse (Figure S8B) appears to be a product of its high seed number (Figure S8C) high harvest index (Figure S8D), and relatively high TGW (Figure S9).

The yield data The cumulative transpiration, taken as the sum of the daily transpiration of each plant throughout the experiment) enabled us to calculate the water use efficiency (WUE) for the 18 lines (Figure 8), which ranged from 0.004 to 0.009 g g^-1^, with Eero having the lowest (0.0045 ± 0.00032 g g^-1^) and Chanell having the highest (0.009 ± 0.0003 g g^-1^). The WUE of Freja and Hydrogen, which displayed during pre-treatment respectively the lowest and highest values for DT (Figure 3A), GSc (Figure 5A), and CPW (Figure 6A), are however, in the mid-range, with Hydrogen being more efficient (0.007 ± 0.0003 g g^-1^) than Freja (0.006 ± 0.0003 g g^-1^).

**Figure 8.**
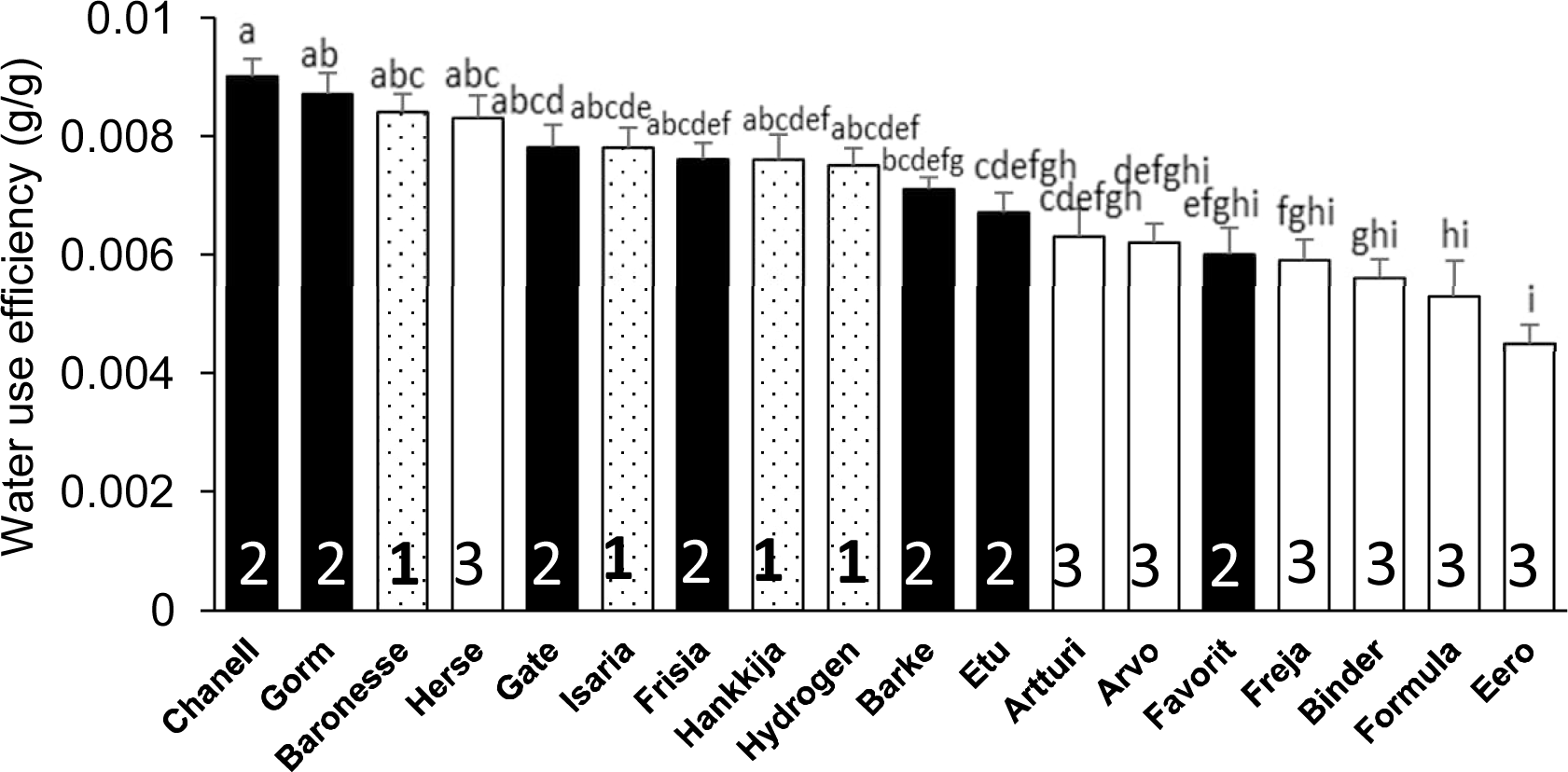
Water use efficiency calculated from dry biomass and cumulative transpiration. Mean ± SE; lowercase letters above the bars are significance groups.

## 4 Discussion

Plants exhibit a wide array of responses and adaptive mechanisms at the morphological, physiological, and molecular levels to drought or water deficit. Nevertheless, the utilization of these mechanisms varies significantly across different plant species or even within genotypes of the same species (Fang and Xiong, 2015), as we have earlier demonstrated for a few chosen varieties of barley (Appiah et al., 2023). Here, we have screened the QPTs of 81 barley genotypes derived from a diversity set spanning 20^th^ Century barley breeding (Tondelli et al., 2013). From the 81 lines, 18 were chosen to represent the range of the QPTs and studied in more detail under well-watered conditions, drought stress, and subsequent recovery.

### 4.1 Performance under well-watered conditions

Our analysis indicated a strong correlation (*r* = 0.95) between vigor (CPW gain per day) and transpiration, those lines (Group 1) with the highest daily transpiration also having the highest vigor. Studies suggest that fractional changes in stomatal conductance lead to changes in transpiration (Drake et al., 2013, Zhang et al., 2019). We found a strong correlation (*r* = 0.94) between daily transpiration and stomatal conductance under well-watered conditions, confirming this (Figure S7B). However, exceptions such as Isaria and Arvo were observed, where the correlation was weak (Figure 3A, 4A). This discrepancy may be related to variations in stomatal density, distribution, size, and number within the cultivated species; our data indicate that Arvo has the highest stomatal density of the lines we have examined (Pereira et al., in prep).

### 4.2 Performance under drought

Across the analyzed lines, we saw wide and significantly different responses to drought. Group 1 lines, particularly Hydrogen, Hankkija 673 and Baronesse, which displayed the highest transpiration (E_max_; Figure 7A), total daily transpiration (DT; Figure 3), canopy stomatal conductance (GSc; Figure 5), daily CPW (Figure 6), and WUE (Figure 7B) when well-watered, shift as a group from a water non-conserving to a water-conserving strategy during drought. Group 1 then displays the lowest DT, GSc, and daily CPW gain. Notably, most Group 1 members reach θ_c_ at a higher SWC than do other lines (Figure 7A), between day 16 to 18 (drought began day 12), aligning with their higher transpiration rates. Gate, a Group 2 member by its behavior under well-watered conditions, clustered with Group 1 in having low DT, GSc, and daily CPW gain under drought.

The lines in Group 3 displayed the opposite behavior as those in Group 1, having low DT, GSc, and daily CPW gain under well-watered conditions but comparatively high values under drought. Among Group 3, cv Freja was the most extreme regarding transpiration under non-limiting and limiting water, and likewise reached θ_c_ at the lowest SWC, on the final day of the drought (day 26). Herse and Artturi, respectively having among the lowest DT under well-water conditions, had together with Freja the highest DT and GSc as well as CPW gain under drought. The three lines also transitioned to θ_c_ at the lowest SWC among the 18 examined in detail. Lines in Group 2, with the exception of Gate as described above, took a middle road regarding transpiration parameters both before and during drought, as well as reaching θ_c_ at intermediate SWC.

### 4.3 Recovery, Resilience, and yield

Following the drought treatment, resumption of a full irrigation regime led to an increase in DT in all lines. Only two line – Hydrogen and Baronesse – displayed simultaneously high initial DT (Figure 3A) as well as high vigor and recovery and good resilience (Table S5, Figure 7A), Only Hydrogen showed both low drought DT (Figure 3B) and high initial GSc (Figure 5A). A similar degree of resilience, which is the degree to which a plant resumes growth, could in principle be obtained with either low, medium or high vigor; the line need only return to its initial state. In fact, we observed that the most resilient lines (Table S5. Figure 7C) were in fact all members of Group 3—Binder, Freja, and Formula—in terms or returning most fully to their initial growth rate, which was in any case low. Freja, Formula and Binder also showed the best recovery of pre-drought DT, likewise initially low; among Group 1, Hydrogen and Baronesse recovered DT best. During rewatering, Isaria, Hydrogen, and Baronesse (all Group 1) returned to the highest rates of vigor (Figure 3C); Isaria showed the best resilience of Group 1.

We examined which of the physiological measures correlated with the final biomass and yield (Figures S8, S9) in pots on the lysimeter. The top five for biomass included three from Group 1 and two from Group 2; while all seven from Group 3 (the anisohydric lines) were found at the bottom of the range. The top of the DT range (Group 1, Figure 3) during pre-treatment and rewatering, as well as for pre-treatment and rewatering CPW vigor (Figure 6A, 6C), showed the best correlation with harvest biomass (Hydrogen, Baronesse, and Isaria) or yield (Hankkija 673). The superior yield of Hankkija 673 despite its mediocre biomass is due to its first-rank harvest index, being a six-row variety, and above average grain weight (Figure S9). Yields in greenhouse pot experiments, with highly constricted soil volumes and root architectures, are not expected to be closely similar to those from field experiments. Nevertheless, on the lysimeter, carbon capture as indicated by CPW and as the tradeoff for DT still correlates with final biomass.

### 4.4 Drought response strategy

The set of lines examined here can be categorized by the two formally contrasting drought strategies, “isohydric”, or water-conserving, and “anisohydric”, or water-non-conserving (Dalal et al., 2017, Moshelion et al., 2015, Sade et al., 2012). Isohydric plants would be expected to limit transpiration and to transition to θ_c_ at a relatively high SWC in order to maintain constant leaf water potential. Anisohydric plants trade a constant hydricity for higher gas exchange and carbon fixation, which would put them at risk in prolonged droughts. Based on their physiological responses, we classify Freja, Artturi, Herse, and Binder as the most anisohydric (all in Group 3), or least drought-responsive, displaying the highest transpiration under drought and reaching the θ_c_ at low SWC (Figure 7A) on the last days of that phase. Group 2 is more isohydric than Group 3, having higher pre-drought DT and a higher θ_c_ (Figure 7A).

In earlier experiments, a single cultivar, RGT Planet, was identified as representing a third drought response type, which may be described as dynamic or plastic (Appiah et al., 2023). This variety (released 2010) is currently the most popular malting barley in Europe, in part due to its consistent and high yields under farm conditions. On the PlantArray system, RGT Planet displayed high transpiration under well-watered conditions, followed by a moderate transpiration decrease under drought, conferring high resilience and, in the pot experiments, high yields that were not significantly reduced by drought. Here, we have established by screening 81 varieties that the dynamic drought strategy is not unique to the relatively new RGT Planet but is found in older lines as well. Group 1, including the “star” lines Hydrogen and Baronesse, displays this dynamic response to drought, transitioning from high-transpiration, water non-conserving to water-conserving behavior, with θ_c_ at high SWC (Figure 7A). While two- and six-row barley generally derive from distinct breeding programs, resulting in genetic structure (Figure S1), the lines analyzed here did not divide categorically by row number into drought response strategies. Neither did drought strategy correlate with year of release: Group 1 spans 1924 (Isaria) to 1999 (Hydrogen).

Ultimately, from the breeder’s and farmer’s perspectives, the optimal drought response strategy is the one that produces the highest sustainable yield and a tolerable annual variation, commensurate with yearly fluctuations in the growth environment -- drought, in this case -- and a particular level of inputs. A high WUE may favor effective growth and ultimately yield when water is limiting; two Group 2 (isohydric) lines, Chanell and Gorm, together with Baronesse of Group 1 (dynamic), showed the highest (Figure 7B). Yield stability despite an early-season drought is expected to benefit from recovery of carbon fixation and thereby growth, as reflected in our experiments by DT and vigor (CPW). For the varieties, growth conditions, and drought treatment here, membership in Group 1, with a dynamic strategy, tended to favor higher biomass. Although they reached θ_c_ at a high SWC, Group 1 members nevertheless lost weight (i.e. dehydrated) most rapidly (Figure 6B) during the drought treatment while displaying a low DT during drought (Figure 3B). Isaria showed the highest rewatering DT and yet reached θ_c_ at a lower SWC, while Hankkija 673 responded at a high SWC, but recovered to a low DT.

We observed a correlation between transpiration (DT, Figure 3; GSc, Figure 5; E_max_, Figure 7A) and SWC at θ_c_ (Figure 7A), with the dynamic lines (Group 1) having both the transpiration measures and θ_c_ high and the anisohydric both low (Group 3). This raises the questions of how and why the high DT, high vigor lines respond quickly to decreases in SWC, but the low DT lines reach θ_c_ only at low SWC, i.e. what controls stomatal aperture as SWC falls, and what the consequences of these water-use behaviors are. Enhanced GSc (such as in Group 1) is correlated with higher demand for root and stem hydraulic conductivity (Kudoyarova et al., 2011); increased demand under restricted hydraulic conductance from drying soil drives down water potential in the water column between root and leaf (Abdalla et al., 2022, Carminati and Javaux, 2020, Yang et al., 2023). Abundant evidence connects stomatal closure to drying signals from roots, with ABA and likely other factors implicated in the signaling (Saradadevi et al., 2017); ABA plays a central role in stomatal closure (Assmann and Jegla, 2016, Hsu et al., 2021). The overall picture therefore suggests that high DT, GSc, E_max_ lines, which are mostly found in Group 1, place high demand on hydraulic conductivity, which becomes limiting, setting off a drought signal to the stomatal guard cells at even moderately reduced SWC. Low conductivity lines, mostly in Group 3, in this model, put more limited demands on conductivity; SWC falls greatly without triggering stomatal closure, a behavior which has been characterized as non-conserving or anisohydric.

The rapid response to falling SWC among Group 1 was associated, particularly for Hydrogen and Baronesse, with the highest degree of recovery. Even though their weight fell rapidly during drought, Isaria, Hydrogen, and Baronesse were the quickest to gain weight during rewatering (Figure 6C; Table S4), while displaying a high DT (Figure 3). We earlier analyzed the gene networks upregulated and downregulated by drought and recovery in the the anisohydric Golden Promise (Paul et al., 2023); autophagy was downregulated during recovery and thereby implicated in drought response. The connection of autophagy to differing degrees of recovery requires further investigation. The results presented here suggest that a dynamic drought response combined with rapid recovery, such as displayed by Group 1 (and elsewhere by RGT Planet), may offer a good ideotype by which to achieve yield stability under droughts of up to two weeks (conditions here). There have been many efforts to model both stomatal conductance (Buckley, 2005, Damour et al., 2010) and the optimization of conductance vs. carbon gain (Joshi et al., 2022, Sperry et al., 2017). Recently, there have also been efforts to use GWAS and genomic selection to identify genetic components of drought tolerance (Dhanagond et al., 2019, Messina et al., 2023, Moualeu-Ngangue et al., 2020, Ningning et al., 2023). Current collaborative work applying nested association-mapping populations, field and precision phenotyping, ideotyping, modeling, and GWAS may allow us to integrate the theoretical and practical frameworks for drought tolerance.

## Author contributions

A.D. carried out the physiological experiments and collected the data. M.P. analyzed the data and wrote the manuscript. M.J. contributed to the physiological experiments and their analysis. M.M. and A.H.S. designed the experiments. A.H.S. helped interpret the data, revised the manuscript, and was project P.I. All authors read and approved the final version.

## Acknowledgements

We acknowledge and thank Anne-Mari Narvanto for excellent technical assistance and Triin Vahisalu for useful discussions on the physiology and molecular biology of drought response.

## Data availability statement

The data that support the findings of this study can be found in the supplementary material of this article or are available from the corresponding author upon reasonable request.

## Supporting information

Figure S1. Principal Components Analysis (PCA) plot of barley diversity set

Figure S2. Plants on the lysimeter platform during 18-line experiment

Figure S3. Daily Transpiration (DT) of the 18 barley lines

Figure S4. Daily transpiration (DT) correlations (*r*) between experimental phases

Figure S5. Correlations among transpirational measures.

Figure S6. Canopy Stomatal conductance (GSc) correlations (*r*) between of the treatments

Figure S7. Correlations for Calculated plant weight (CPW) gain per day between treatments

Figure S8. Yield estimates for the 18 barley lines from the second screening

Figure S9. Thousand Grain Weight for the 18 barley lines

Table S1. 81 spring barley lines for first screening

Table S2. Characteristics selected 18 spring barley lines

Table S3. TR_max_ and θ_c_ from 81-line screen

Table S4. E_max_ and θ_c_ from 18-line

Table S5. Summary of 18-line physiological strategy and performance

