## Supplemental figures and tables for "Precision phenotyping of a barley diversity set reveals distinct drought response strategies"

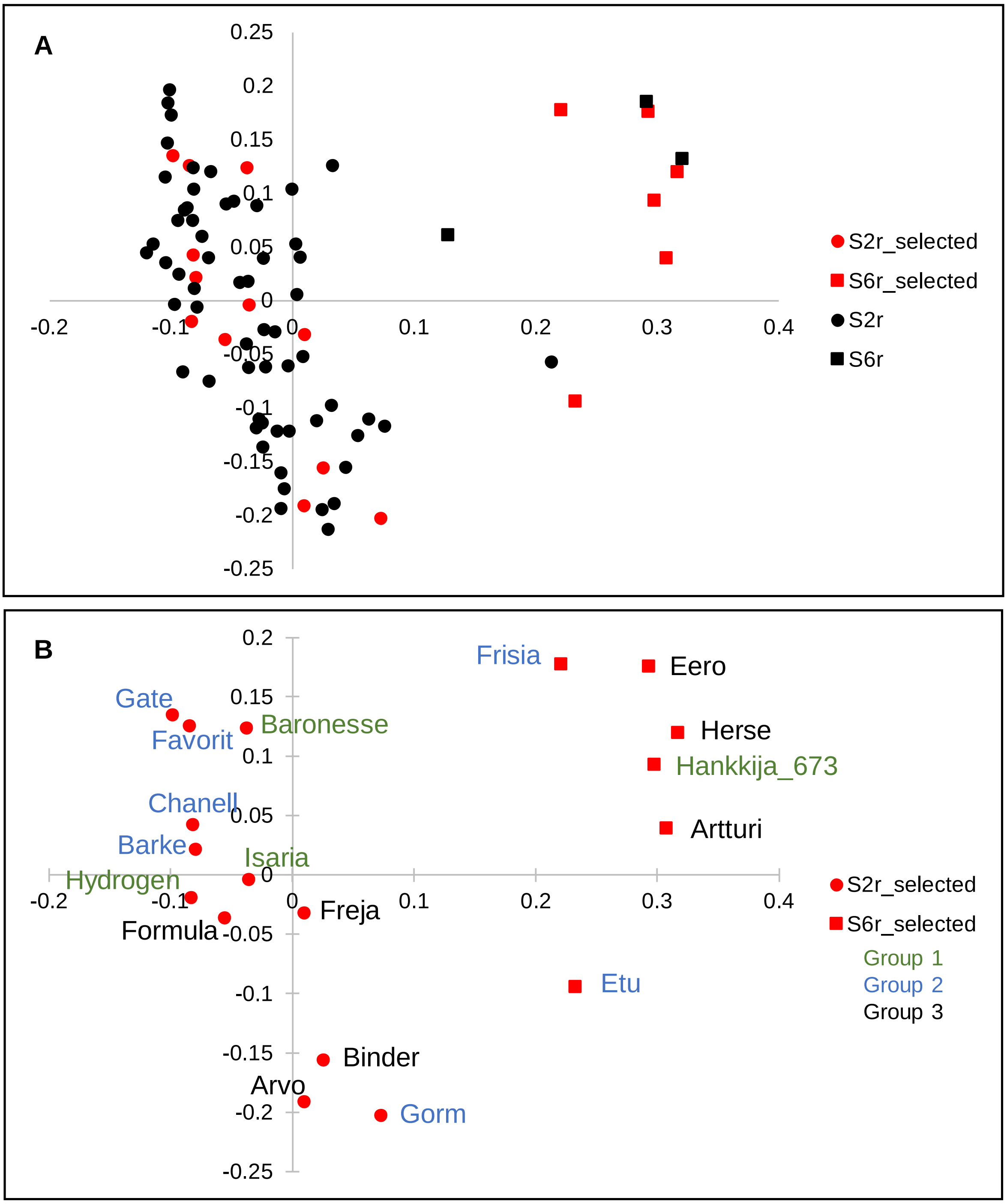

Figure S1. Principal component analysis (PCA) of genome-wide single-nucleotide polymorphism (SNP) data for barley lines under analysis. (A) 81-line set (B) 18-line subset (red). Circles represent two-row, squares six-row, cultivars. Physiological groups are labeled as Group 1 (green), Group 2 (blue) Group 3 (black).

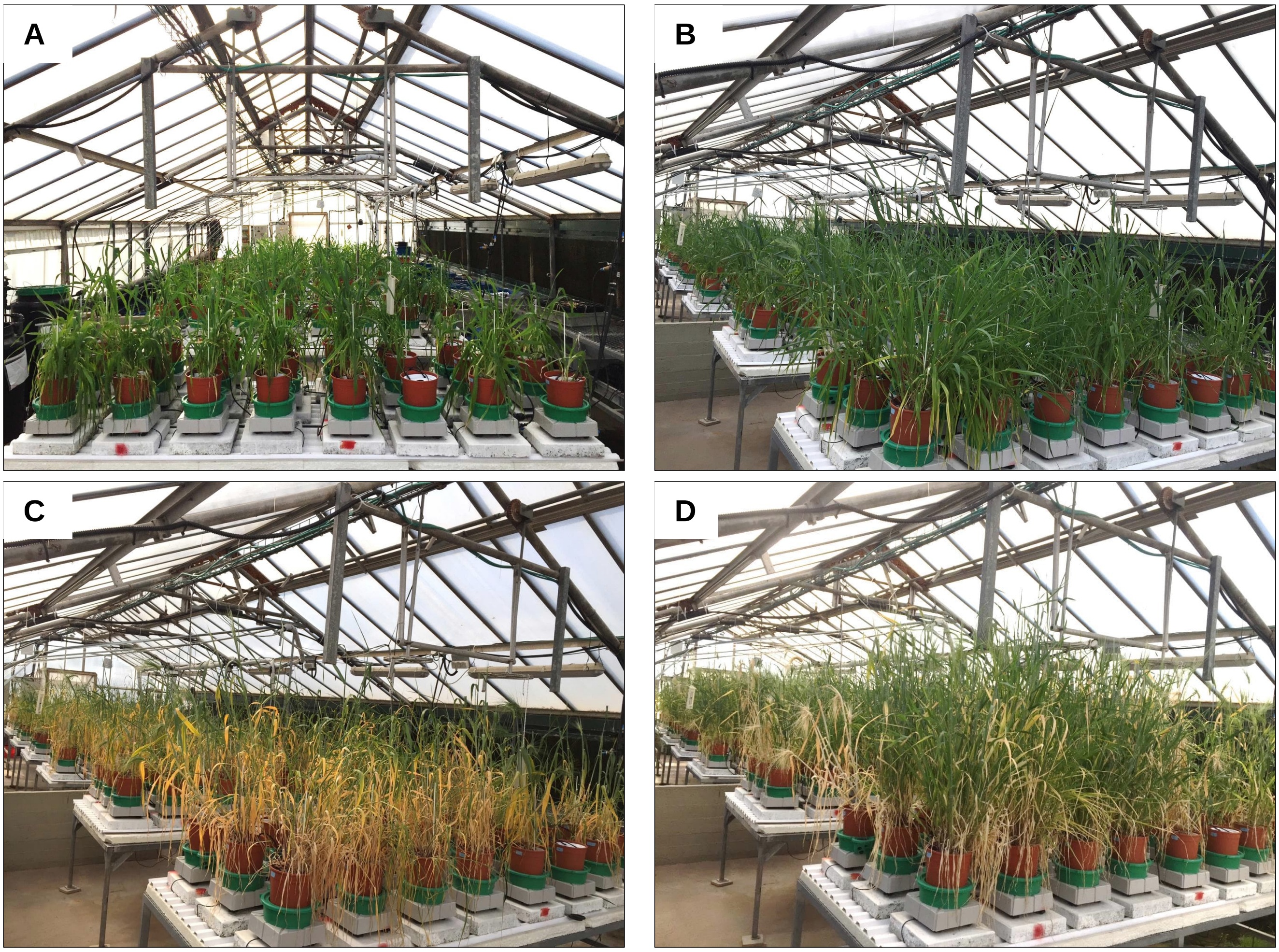

Figure S2. Plants on the lysimeter platform during 18-line experiment. (A) 98-day-old plants, before the pre-treatment. (B) 110 days, during pre-treatment, (C) 125 days, immediately post-drought, (D) 158 days.

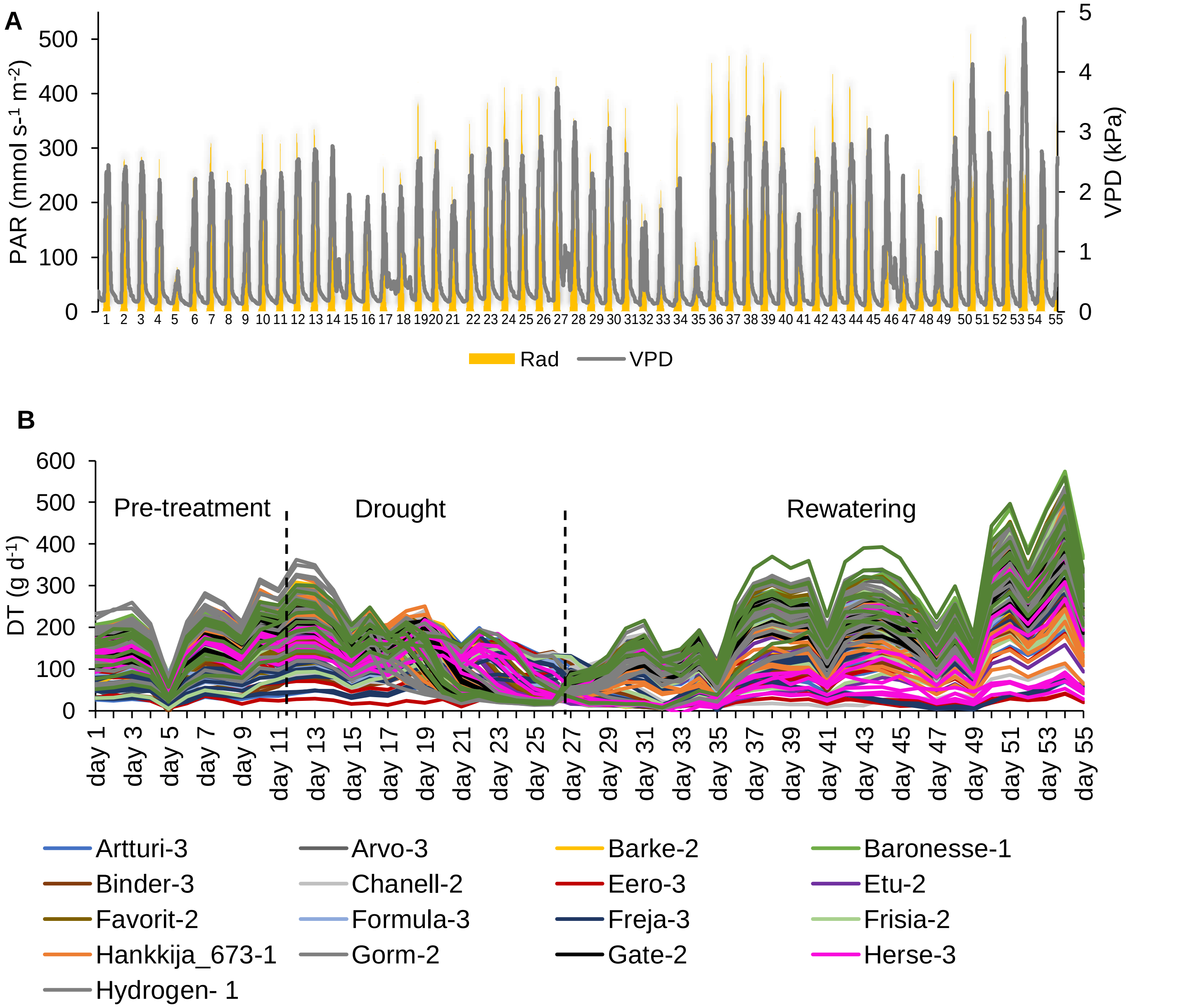

Figure S3. Daily Transpiration (DT) of the 18 barley lines with five to eight biological replicates (139 plants) grown on the lysimeter system for second screening, subjected to full irrigation for 11 days, followed by a 15 days of drought treatment and then rewatering for 29 days. (A) Daily vapor pressure deficit (VPD) and Photosynthetic active radiation (PAR) during 55 consecutive days of the experiment. (B) DT in response to the soil-atmosphere water gradient during 55 consecutive days of the experiment. Each line represents one single plant from the 18 barley accessions. Dashed vertical black line at day 12 and day 26 shows the change in treatment.

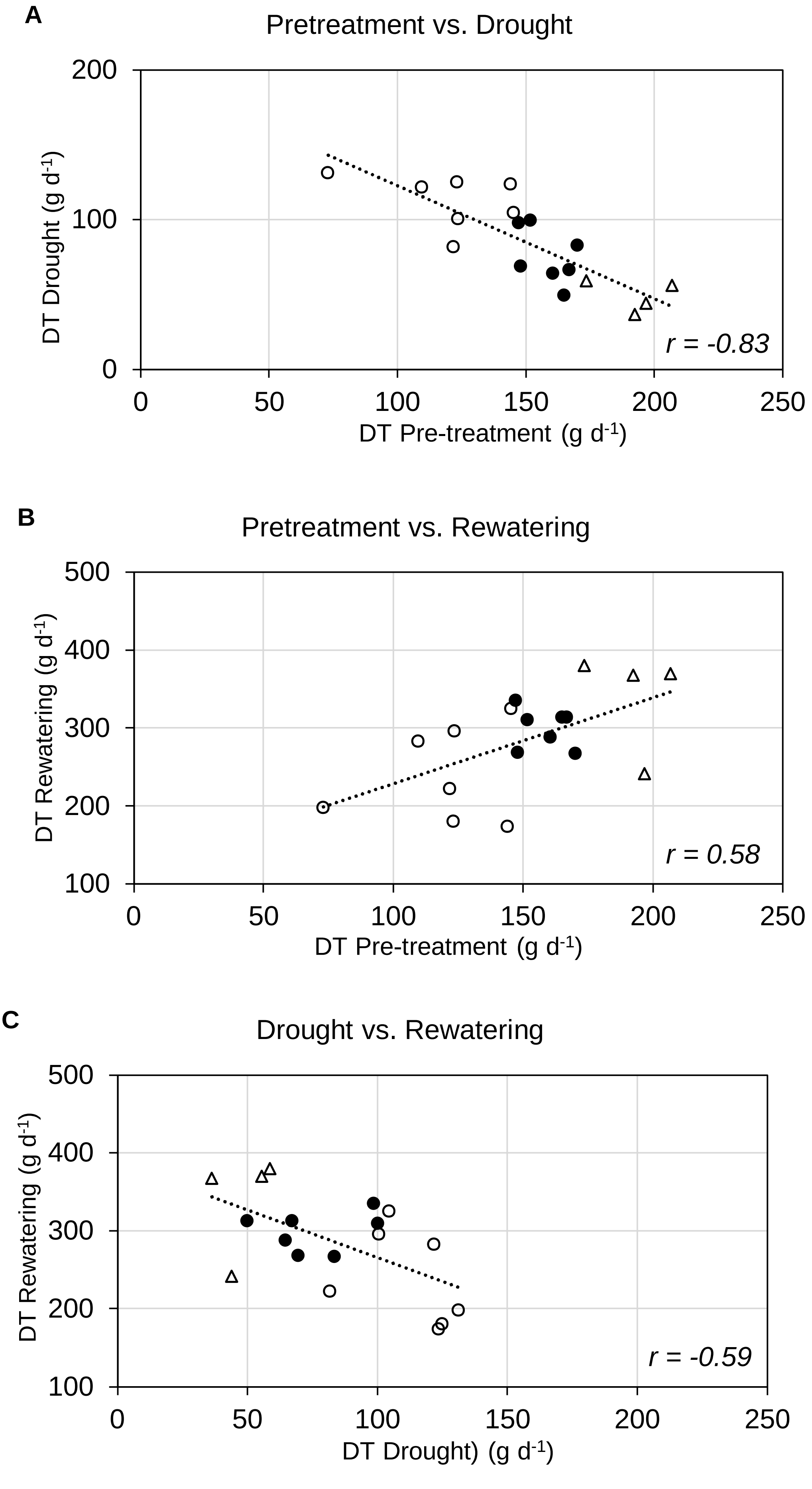

Figure S4: Daily transpiration (DT) correlations (*r*) between experimental phases. Scatter plot between (A) Pre-treatment and Drought (B) Pre-treatment and Rewatering, and (C) Drought and Rewatering. Group 1 (open triangles) shows the highest DT, Group 3 (open circles) show the lowest DT and Group 2 (filled circles) intermediate DT in pre-treatment.

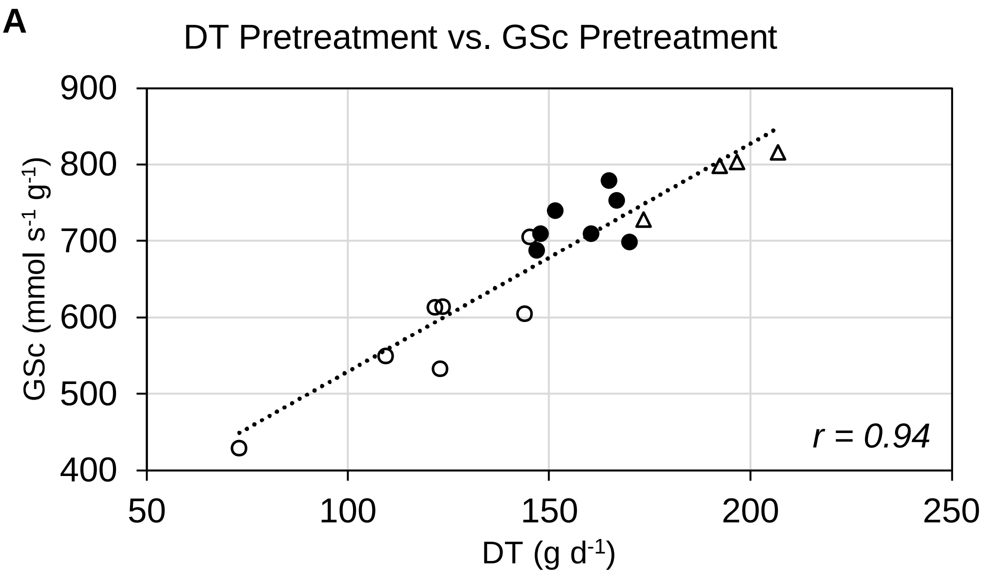

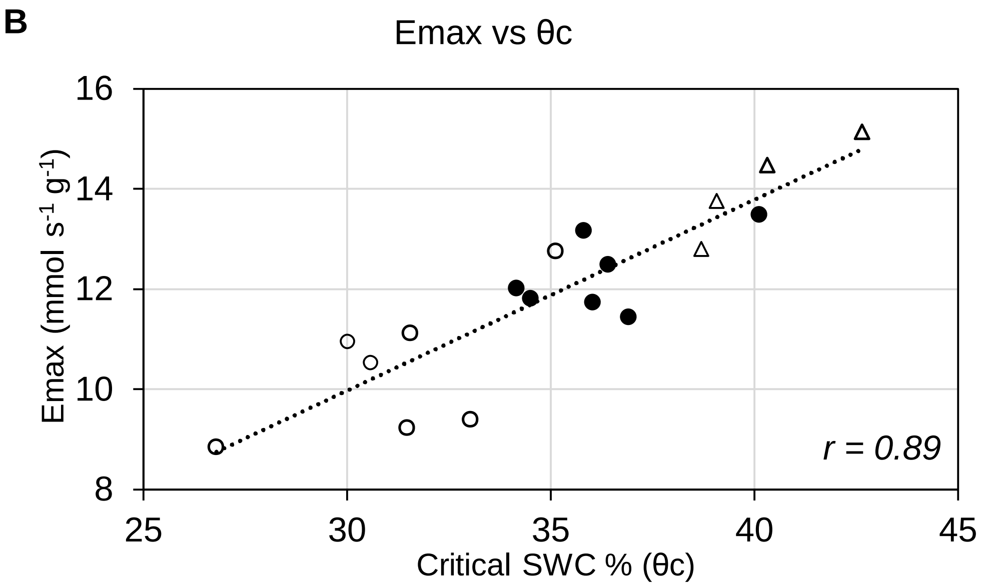

Figure. S5. Correlations among transpirational measures. (A) Correlation between DT (Daily transpiration) and GSc (Canopy Stomatal conductance) during pre-treatment. (B) Correlation between Emax and the theta critical SWC (θc). Group 1, open triangles; Group 2, filled circles; Group 3, open circles.

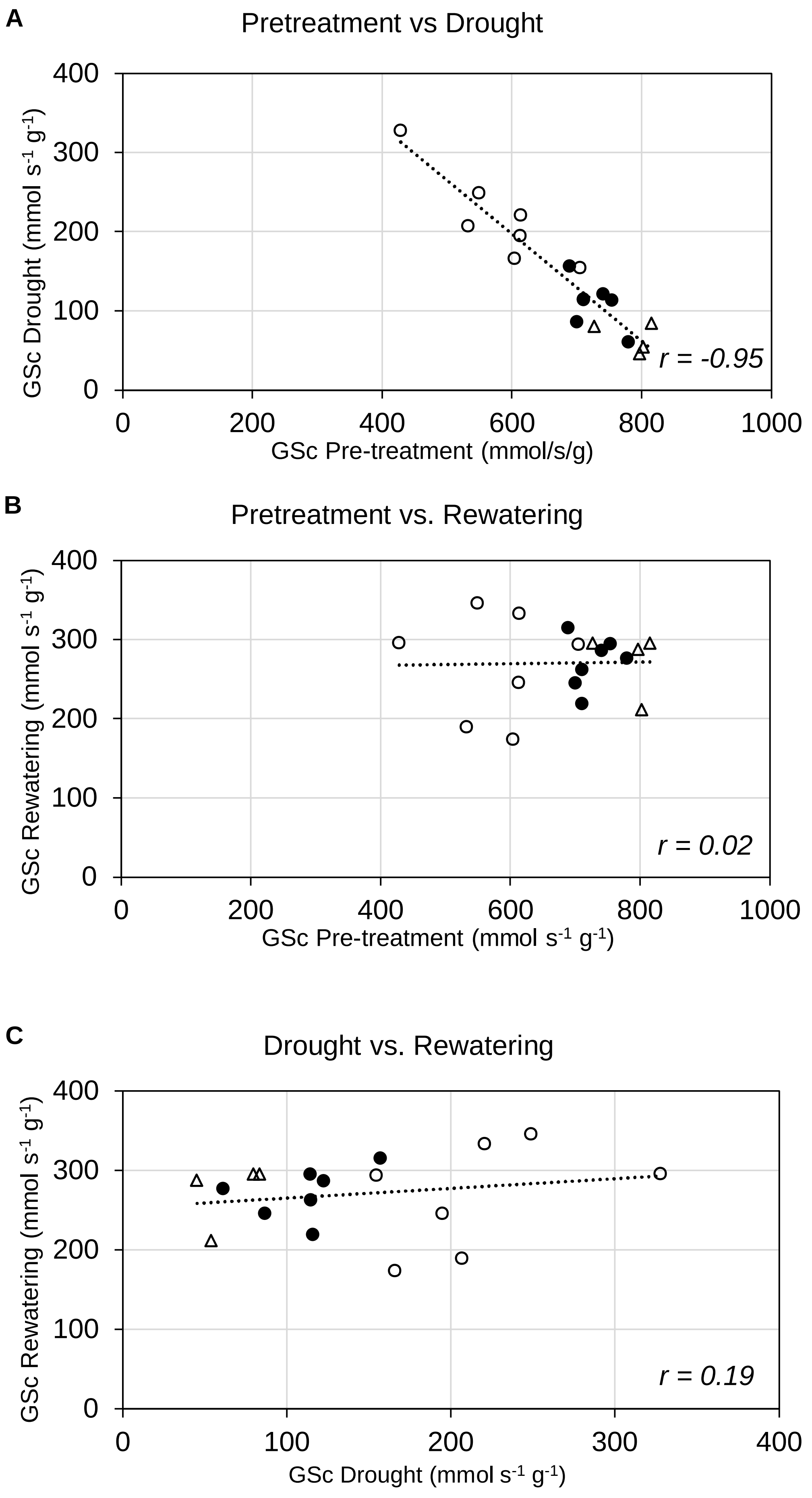

Figure S6: Canopy Stomatal conductance (GSc) correlations (*r*) between of the treatments. Scatter plot between (A) Pre-treatment and Drought (B) Pre-treatment and Rewatering, and (C) Drought and Rewatering. Group 1 (open triangles), Group 2 (filled circles), Group 3 (open circles).

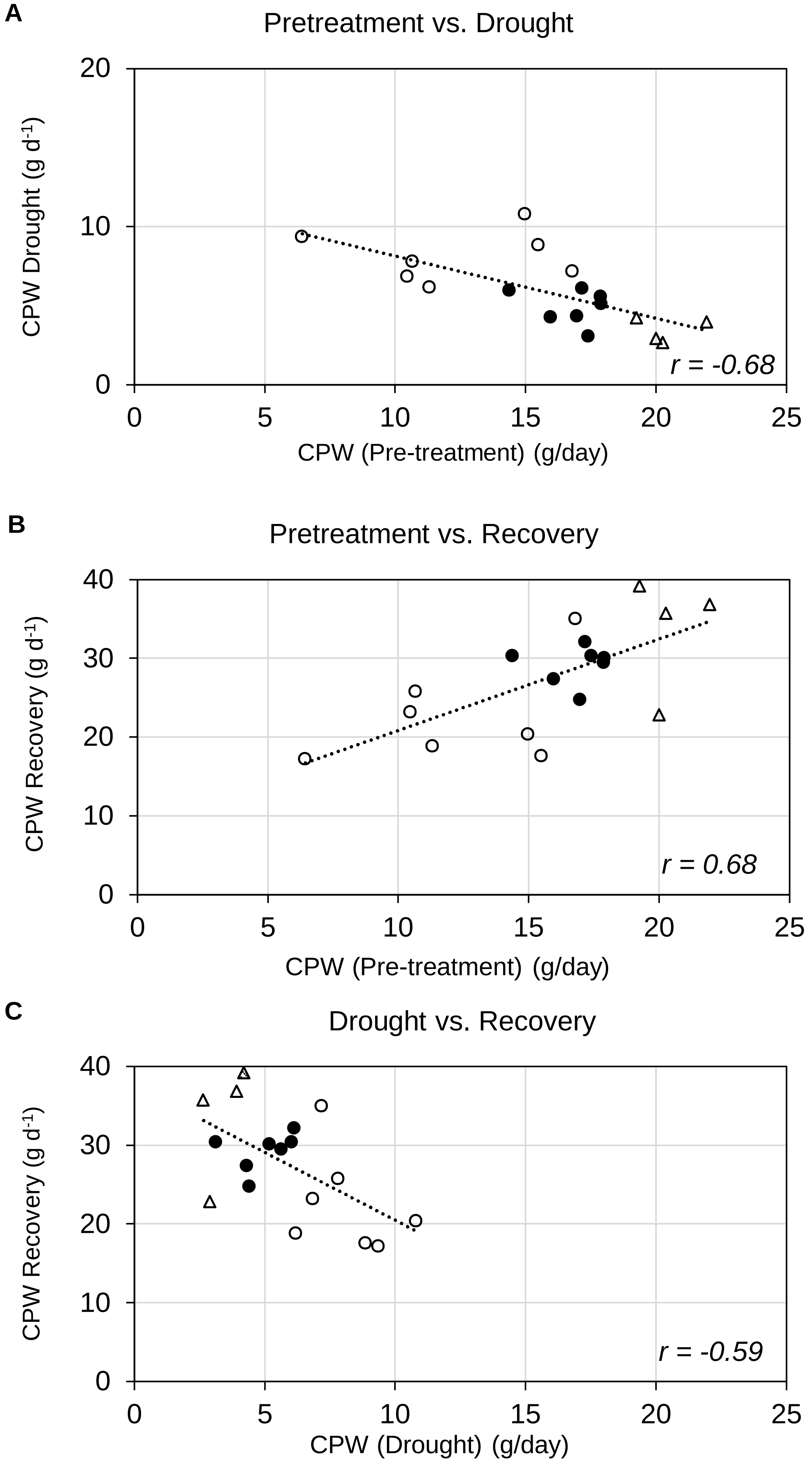

Figure S7: Correlations for Calculated plant weight (CPW) gain per day between treatments. Scatter Plot between (A) Pre-treatment and Drought. (B) Pre-treatment and Recovery. (C) Drought and Recovery. Group 1, hatched circles; Group 2, filled circles; Group 3, open circles.

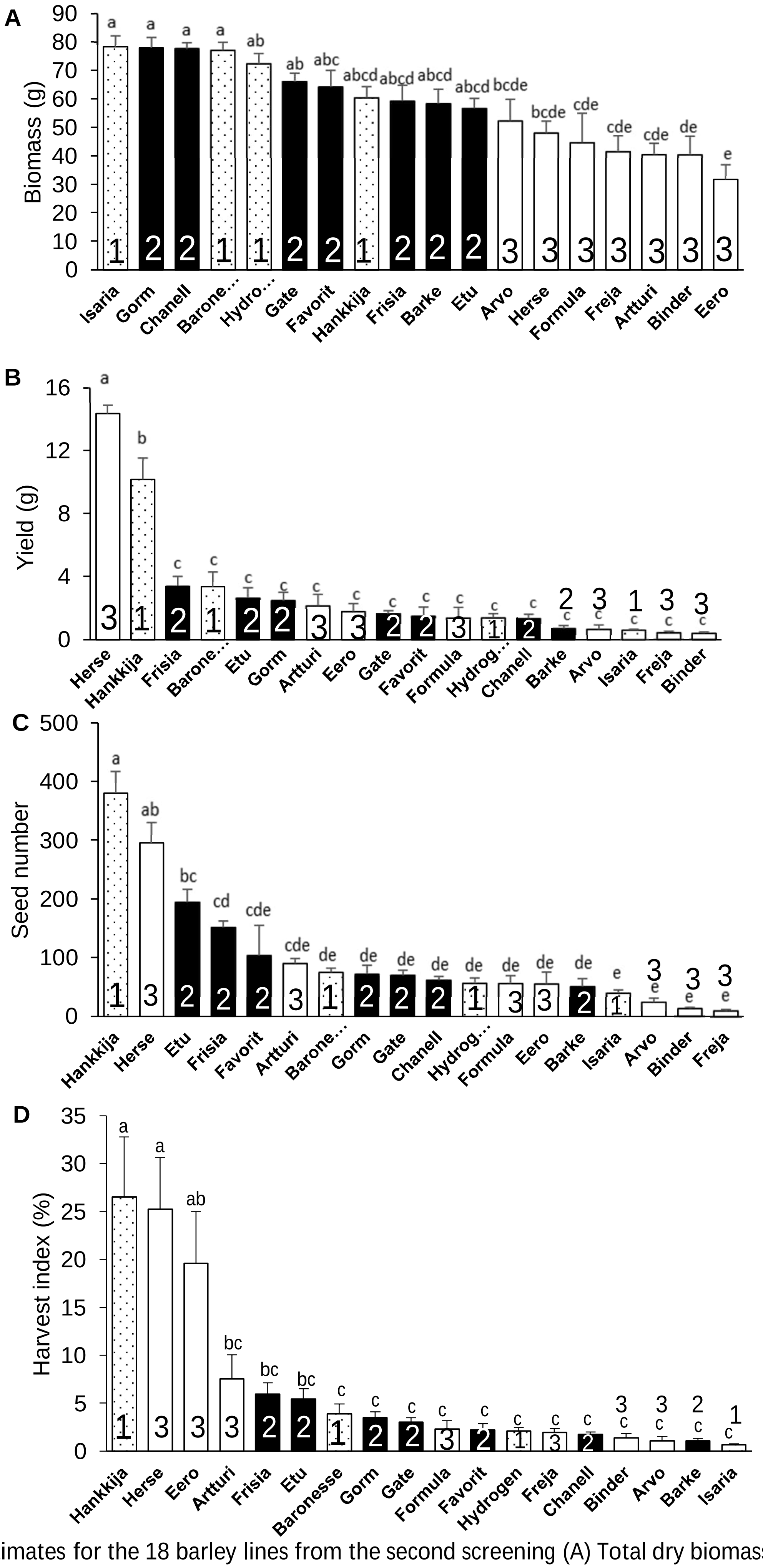

Figure S8: Yield estimates for the 18 barley lines from the second screening (A) Total dry biomass (weight of husk, seed, and shoot). (B) Yield per line. (C) Seed number. (D) Harvest index (%). Mean ± SE; lowercase letters indicate significance groups.

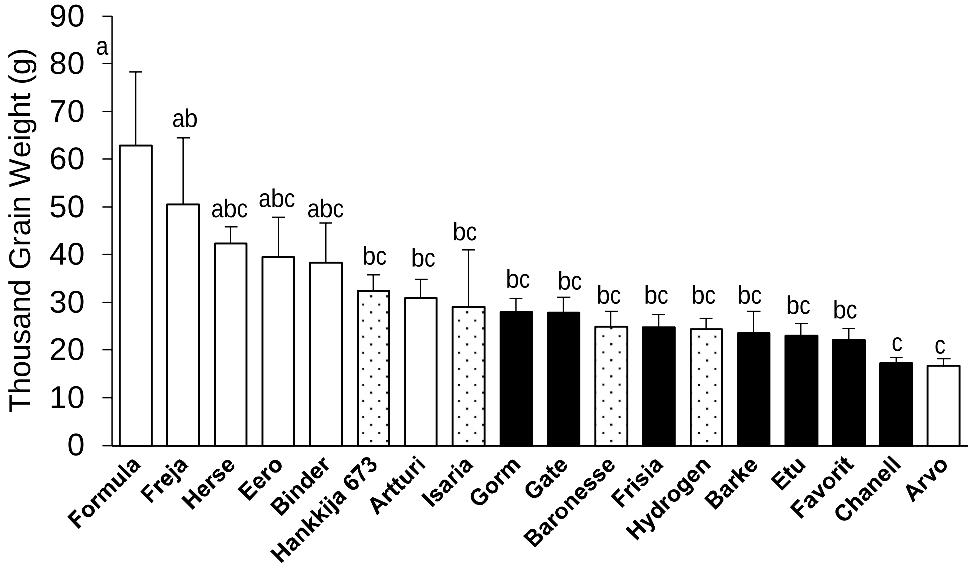

Figure S9 Thousand Grain Weight for the 18 barley lines. Mean ± SE; lowercase letters indicate significance groups.

Table S1. 81 spring barley lines for first screening

| Accession | Code | Name | Rows | Release | Country | Pedigree | Breeder |
|---|---|---|---|---|---|---|---|
| 126 | 2009 | Aapo | 2 | 1975 | Finland | Carlsberg x Riegel | Hankkija Plant Breeding Institute |
| 128 | 2010 | Abava | 2 | 1978 | Latvia | Mari/Elsa//Domen | Stende Plant Breeding Station |
| 129 | 2011 | Agneta | 6 | 1981 | Sweden | Asa x Frisia x EddaII x Monte Christo | Svalöf |
| 131 | 2012 | Akcent | 2 | 1992 | Czech Republic | Salome/EP 79 | Selgen Stupice |
| 132 | 2013 | Akka | 2 | 1970 | Sweden | (Arla)6 x Monte Cristo | Weibull |
| 135 | 2015 | Alis | 2 | 1985 | Denmark | Triumpf x Rosie Abed | Abed |
| 136 | 2016 | Alliot | 2 | 1999 | Denmark | Chariot x Alexis | Pajbjergfonden |
| 137 | 2018 | Anni | 2 | 1980 | Estonia | Lola x Liisa | Jogeva Plant Breeding Institute |
| 142 | 2019 | Ansis | 2 | 1996 | Latvia | Jarek/Taifun | Stende Plant Breeding Station |
| 143 | 2020 | Apex | 2 | 1984 | Netherlands | Aramir x F1[CEB 6721 x (Julia3 x Volla x L100)] | Cebeco |
| 144 | 2021 | Aramir | 2 | 1974 | Netherlands | Volla x Emir | Cebeco |
| 148 | 2024 | Artturi | 6 | 1992 | Finland | Arra x Nord | Boreal Plant Breeding Ltd |
| 149 | 2025 | Arvo | 2 | 1966 | Finland | Balder x Helmi | Agricultural Experiment Station of Finland, Jokioinen |
| 153 | 2027 | Athos | 2 | 1975 | France | 207*Emir | Desprez |
| 154 | 2028 | Atlas | 2 | 1976 | Czech Republic | M- SS 55/Diamant | Selgen Stupice |
| 155 | 2029 | Atribut | 2 | 1996 | Czech Republic | KM V 3-83/BR 2174 | Selgen Stupice |
| 158 | 2030 | Balder J | 2 | 1964 | Sweden | Balder X-Ray Mutant | Weibull |
| 159 | 2031 | Balga | 2 | 1990 | Latvia | Gunilla/KM 1192 | APP Valsts Priekulu laukaugu selekcijas instituts |
| 160 | 2032 | Barabas | 2 | 2005 | Denmark | SJ 970621 x (Lux x Annabell) | Sejet |
| 162 | 2033 | Barke | 2 | 1996 | Germany | Libelle x Alexis | Breun |
| 163 | 2034 | Baronesse | 2 | 1989 | Germany | ((343/6 x V34/6) x J -427) x (Oriol x LBW6153 P40) | Nordsaat |
| 164 | 2035 | Beatrix | 2 | 2004 | Germany | Viskosa x Pasadena | Nordsaat |
| 165 | 2036 | Berenice | 2 | 1972 | France | Union*((Bordia*Kenia)*Frisia) | INRA |
| 166 | 2037 | Binder | 2 | 1916 | Denmark | HOR3684/76 ABED selection in Hanna | Abed |
| 168 | 2039 | Birka | 2 | 1981 | Sweden | Baladi16 x (Rika x ((Tellus x (Monte_Christo x Tellus MD)) | Weibull |
| 169 | 2040 | Blenheim | 2 | 1992 | United Kingdom | Triumph*Egmont | PBI |
| 173 | 2043 | Braemar | 2 | 2000 | United Kingdom | NFC 5563 x NFC 94.20 | New Farm Crops (NFC) |
| 174 | 2044 | Brazil | 2 | 2001 | France | Trebon x Cooper | Momont |
| 176 | 2045 | Britta A | 2 | 1964 | Sweden | (Binder*Opal)*(Balder*Kenia) | Lantmännen SW Seed |
| 180 | 2046 | Caja | 2 | 1980 | Denmark | PF-M-13 x PF 62-6/6-4 | Pajbjerg |
| 187 | 2048 | Carlsberg II | 2 | 1947 | Denmark | Prentice x Maja | Carlsberg |
| 190 | 2050 | Ceylon | 2 | 2003 | Netherlands | Portia x Amber | Cebeco |
| 193 | 2051 | Chanell | 2 | 2006 | Denmark | Barke x Ca 500201 | Carlsberg |
| 194 | 2052 | Chariot | 2 | 1992 | United Kingdom | Dera*(Carnival*Atem) | PBI |
| 200 | 2053 | Claret | 2 | 1980 | United Kingdom | ((Proctor*HP 5466)*Armelle)*Abacus | Nickerson |
| 201 | 2054 | Class | 2 | 2003 | United Kingdom | Prestige x Optic | PBI |
| 205 | 2056 | Cooper | 2 | 1994 | United Kingdom | (Corniche*Force)*Troop | New Farm Crops (NFC) |
| 206 | 2057 | Corgi | 2 | 1983 | United Kingdom | Triumph*15533 Co | Welsh Pl Breed Stn |
| 210 | 2059 | Dandy | 2 | 1988 | United Kingdom | Egmont*Atem | Welsh Pl Breed Stn |
| 209 | 2060 | Croydon | 2 | 1982 | Sweden | welam x tellus $M_1D$ | Weibull |
| 208 | 2061 | Cristalia | 2 | 2004 | United Kingdom | Ortoli x Brise | Syngenta |
| 211 | 2062 | Danuta | 2 | 2000 | Germany | 90014DH [Krona] x (Salome x Maresi) | Nordsaat |
| 212 | 2063 | Deba Abed | 2 | 1965 | Denmark | Abed Denso*Weihenstephaner MR 2 | Abed |
| 213 | 2064 | Delta | 2 | 1959 | Netherlands | Tyra*Claret OR Kenia*H.laevigatum*Gull | Cebeco |
| 214 | 2065 | Derkado | 2 | 1992 | Germany | Lada*Salome | Hadmersleben |
| 215 | 2066 | Dialog | 2 | 2000 | Denmark | Otira x (Ferment x Mentor) | Sejet |
| 216 | 2067 | Diamant | 2 | 1965 | Czech Republic | Valticky X-ray Mutant | OGZ |
| 217 | 2068 | Digersano | 2 | 1991 | Italy | Mari-Coho x Sul-Mackta | Cermis/ENEA |
| 223 | 2070 | Drost | 2 | 1954 | Denmark | Maja x Kenia | Pajbjerg |
| 224 | 2071 | Druvis | 6 | 1999 | Latvia | Dobrij/HVS 115440 | Stende Plant Breeding Station |
| 226 | 2072 | Edda | 6 | 1949 | Sweden | Vega x Asplund | Svalöf |

| DETAILS | |
|---|---|
| Year | 1915-2006 |
| 2- row | 72 |
| 6- row | 9 |
| Breeders | 38 |
| Austria | 1 |
| Czech Republic | 9 |
| Denmark | 14 |
| Estonia | 2 |
| Finland | 7 |
| France | 3 |
| Germany | 9 |
| Italy | 1 |
| Latvia | 5 |
| Netherlands | 5 |
| Norway | 1 |
| Slovak Republic | 1 |
| Sweden | 12 |
| United Kingdom | 11 |
| 14 countries | 81 |

| | | | | | | | |
|---|---|---|---|---|---|---|---|
| 227 | 2073 | Eero | 6 | 1975 | Finland | Mari 2r x Otra | Hankkija Plant Breeding Institute |
| 228 | 2074 | Egmont | 2 | 1980 | United Kingdom | (Maris Yak*W 1001)*Vada | PBI |
| 229 | 2075 | Elantra | 2 | 1998 | Denmark | Caminant x Heron | Sejet |
| 230 | 2076 | Elo | 2 | 1989 | Estonia | Triumph x Lofa | Jogeva Plant Breeding Institute |
| 231 | 2077 | Emir | 2 | 1962 | Netherlands | Delta*(Agio*(Kenia)2*Arabian Variety) | Cebeco |
| 234 | 2079 | Etu | 6 | 1970 | Finland | Bonus M x Varde | Boreal Plant Breeding Ltd |
| 235 | 2080 | Eunova | 2 | 2000 | Austria | (Serva x ML502) x CF 79 | Probstdorf |
| 237 | 2081 | Famin | 2 | 1996 | Czech Republic | Akcent/CE 597 | Hrubcice |
| 238 | 2082 | Favorit | 2 | 1973 | Czech Republic | Diamant/F.Union | Hrubcice |
| 239 | 2083 | Felicitas | 2 | 2002 | Germany | (Baronesse x Meltan) x Krona) | Breun |
| 240 | 2084 | Formula | 2 | 1987 | Sweden | Triumph x A 11 3109 | Weibull |
| 241 | 2085 | Forum | 2 | 1993 | Czech Republic | H 387-75/Horpatsi Ketscoros//044-78 | HYBRITECH |
| 242 | 2086 | Freja | 2 | 1941 | Sweden | Victory X Opal | Svalöf |
| 243 | 2087 | Frisia | 6 | 1955 | Germany | (Granat*Pirthgjarn)*(Eckendorfer WG*Kalckreuthen WG) | Breustedt |
| 245 | 2088 | Galan | 2 | 1990 | Slovak Republic | Complex hybrid. K 2567/HE 1428 | Sladkovicovo |
| 267 | 2089 | Helmi | 2 | 1942 | Finland | Binder X Pikkio | na |
| 268 | 2090 | Heris | 2 | 1998 | Czech Republic | HE 4431/CE 431 | Hrubcice |
| 51 | 2091 | Harry | 2 | 1978 | Sweden | Maythorpe Gamma-Ray Mutant | Weibull |
| 260 | 2092 | Hanna | 2 | 1992 | Sweden | (Armelle*Lud)*Luke | Weibull |
| 259 | 2093 | Hankkija_673 | 6 | 1973 | Finland | (Herta 8 x Byg 191 x Ingrid x Minerva) x Kristina | Hankkija Plant Breeding Institute |
| 258 | 2094 | Hanka | 2 | 1997 | Germany | Swedish (Gotland) Land Variety | Semundo |
| 257 | 2095 | Hana | 2 | 1973 | Czech Republic | Diamant/Alsa | OGZ |
| 255 | 2096 | Gull | 2 | 1915 | Sweden | Swedish Land Variety | Svalöf |
| 254 | 2097 | Gorm | 2 | 1981 | Denmark | Otra x Paavo | Sejet |
| 253 | 2098 | Golf | 2 | 1983 | United Kingdom | [(monte cristo x 5690) x $5793^{2}$] x $5853^{3}$ | Nickerson |
| 250 | 2100 | Gizmo | 2 | 2006 | Denmark | Prestige x Ca 800602 | Carlsberg |
| 247 | 2101 | Gate | 2 | 1995 | Latvia | Emir/2*Nadja//HE-497/Hadmersleben 70197/70 | Priekuli |
| 269 | 2102 | Herse | 6 | 1939 | Norway | Asplund x Maskin | Vollebekk |
| 271 | 2103 | Hydrogen | 2 | 1999 | Denmark | (Alis x Digger) x Derkado | Nordic seed |
| 283 | 2104 | Isaria | 2 | 1924 | Germany | Bavaria x Danubia | Ackermann |

Table S2. Characteristics selected 18 spring barley lines

| Accession | Code | Name | Rows | Release | Country | Pedigree | Breeder |
|---|---|---|---|---|---|---|---|
| 148 | 2024 | Artturi | 6 | 1992 | Finland | Arra x Nord | Boreal Plant Breeding Ltd |
| 227 | 2073 | Eero | 6 | 1975 | Finland | Mari 2r x Otra | Hankkija Plant Breeding Institute |
| 243 | 2087 | Frisia | 6 | 1955 | Germany | (Granat*Pirthgjarn)*(Eckendorfer WG*Kalckreuthen \ | Breustedt |
| 193 | 2051 | Chanell | 2 | 2006 | Denmark | Barke x Ca 500201 | Carlsberg |
| 234 | 2079 | Etu | 6 | 1970 | Finland | Bonus M x Varde | Boreal Plant Breeding Ltd |
| 238 | 2082 | Favorit | 2 | 1973 | Czech Re | Diamant/F.Union | Hrubcice |
| 259 | 2093 | Hankkija_673 | 6 | 1973 | Finland | (Herta 8 x Byg 191 x Ingrid x Minerva) x Kristina | Hankkija Plant Breeding Institute |
| 269 | 2102 | Herse | 6 | 1939 | Norway | Asplund x Maskin | Vollebekk |
| 242 | 2086 | Freja | 2 | 1941 | Sweden | Victory X Opal | Svalöf |
| 254 | 2097 | Gorm | 2 | 1981 | Denmark | Otra x Paavo | Sejet |
| 247 | 2101 | Gate | 2 | 1995 | Latvia | Emir/2*Nadja//HE-497/Hadmersleben 70197/70 | Priekuli |
| 271 | 2103 | Hydrogen | 2 | 1999 | Denmark | (Alis x Digger) x Derkado | Nordic Seed |
| 283 | 2104 | Isaria | 2 | 1939 | Germany | Bavaria x Danubia | Ackermann |
| 149 | 2025 | Arvo | 2 | 1966 | Finland | Balder x Helmi | Agricultural Experiment Station of Finland, Jokioinen |
| 162 | 2033 | Barke | 2 | 1996 | Germany | Libelle x Alexis | Breun |
| 163 | 2034 | Baronesse | 2 | 1989 | Germany | ((343/6 x V34/6) x J -427) x (Oriol x LBW6153 P40) | Nordsaat |
| 166 | 2037 | Binder | 2 | 1916 | Denmark | HOR3684/76 ABED selection in Hanna | Abed |
| 240 | 2084 | Formula | 2 | 1987 | Sweden | Triumph x A 11 3109 | Weibull |

| DETAILS | |
|---|---|
| 2- rows | 12 |
| 6- rows | 6 |
| Year | 1916- 2006 |
| Breeders | 16 |
| Czech Repu | 1 |
| Denmark | 4 |
| Finland | 5 |
| Germany | 4 |
| Latvia | 1 |
| Norway | 1 |
| Sweden | 2 |
| 7 countries | 18 |

Table S3. Maximum transpiration rate TRmax uder well-watered conditions; critical SWC **θ**c during drought stress from 81-line screen

| Group | Code | Name | TRmax | **θ**c |
|---|---|---|---|---|
| A | 2024 | Artturi | 10.42 | 35.59 |
| A | 2087 | Frisia | 11.48 | 29.86 |
| A | 2073 | Eero | 12.68 | 30.22 |
| B | 2102 | Herse | 13.95 | 18.06 |
| B | 2051 | Chanell | 14.02 | 25.60 |
| B | 2079 | Etu | 15.79 | 25.18 |
| B | 2093 | Hankkija_673 | 15.88 | 16.83 |
| B | 2082 | Favorit | 16.87 | 23.78 |
| C | 2086 | Freja | 18.34 | 28.97 |
| C | 2104 | Isaria | 18.51 | 27.83 |
| C | 2103 | Hydrogen | 18.79 | 28.01 |
| C | 2101 | Gate | 19.02 | 27.96 |
| C | 2097 | Gorm | 20.22 | 29.40 |
| D | 2025 | Arvo | 22.51 | 30.00 |
| D | 2037 | Binder | 26.20 | 30.00 |
| D | 2084 | Formula | 27.18 | 31.41 |
| D | 2033 | Barke | 28.01 | 30.00 |
| D | 2034 | Baronesse | 29.18 | 30.00 |

Table S4. Maximum transpiration (Emax) under well-watered conditions; critical SWC **θ**c during drought stress, 18 lines

| Group | Code | Name | **θ**c | Std. Error o | Emax | Std. Error of Mean |
|---|---|---|---|---|---|---|
| Group 1 | 2103 | Hydrogen | 42.6533 | 2.918 | 15.1384 | 1.29488 |
| | 2093 | Hankkija_673 | 40.3231 | 2.27564 | 14.4747 | 0.94255 |
| | 2034 | Baronesse | 39.087 | 1.97292 | 13.7502 | 0.50675 |
| | 2104 | Isaria | 38.7049 | 1.50548 | 12.7957 | 0.58368 |
| Group 2 | 2033 | Barke | 40.1149 | 2.31025 | 13.4909 | 0.62889 |
| | 2097 | Gorm | 35.8127 | 2.43632 | 13.1754 | 0.60869 |
| | 2101 | Gate | 36.4074 | 1.11428 | 12.5033 | 0.21707 |
| | 2051 | Chanell | 34.1508 | 1.42127 | 12.0246 | 0.55855 |
| | 2082 | Favorit | 34.5064 | 2.23584 | 11.8246 | 0.57105 |
| | 2087 | Frisia | 36.0204 | 2.58709 | 11.7394 | 0.90145 |
| | 2079 | Etu | 36.9027 | 0.52283 | 11.4477 | 0.57186 |
| Group 3 | 2025 | Arvo | 35.1197 | 2.10051 | 12.7548 | 0.59304 |
| | 2037 | Binder | 31.5549 | 2.24836 | 11.1248 | 0.52813 |
| | 2084 | Formula | 30.0064 | 3.07542 | 10.952 | 1.87731 |
| | 2073 | Eero | 30.5805 | 2.18401 | 10.5412 | 1.57636 |
| | 2102 | Herse | 33.0353 | 1.79695 | 9.3969 | 0.64774 |
| | 2024 | Artturi | 31.4731 | 2.30717 | 9.2269 | 0.70764 |
| | 2086 | Freja | 26.7912 | 1.23907 | 8.8455 | 0.78473 |

Table S5. Summary of 18-line physiological strategy and performance

| Lines | 81-line grouping | 18 line grouping | Drought Strategy | Vigor | Resilience | Recovery |
|---|---|---|---|---|---|---|
| Hankkija_673 | B | 1 | dynamic | high | low | medium |
| Isaria | C | 1 | dynamic | high | medium | medium |
| Hydrogen | C | 1 | dynamic | high | medium | high |
| Baronesse | D | 1 | dynamic | high | medium | high |
| Frisia | A | 2 | isohydric | medium | medium | medium |
| Etu | B | 2 | isohydric | medium | low | low |
| Favorit | B | 2 | isohydric | low | medium | medium |
| Chanell | B | 2 | isohydric | medium | medium | low |
| Gate | C | 2 | isohydric | medium | medium | medium |
| Gorm | C | 2 | isohydric | medium | medium | high |
| Barke | D | 2 | isohydric | medium | medium | medium |
| Eero | A | 3 | anisohydric | low | medium | medium |
| Artturi | A | 3 | anisohydric | medium | low | medium |
| Herse | B | 3 | anisohydric | medium | low | low |
| Freja | C | 3 | anisohydric | low | high | high |
| Arvo | D | 3 | anisohydric | medium | medium | medium |
| Binder | D | 3 | anisohydric | low | high | high |
| Formula | D | 3 | anisohydric | low | high | high |
